# DNA methylation stochasticity is linked to transcriptional variability and identifies convergent epigenetic disruption across genetically-defined subtypes of AML

**DOI:** 10.1101/2024.10.26.620422

**Authors:** Eleanor Hilgart, Weiqiang Zhou, Eduardo Martinez-Montes, Adrian Idrizi, Rakel Tryggvadottir, Lukasz P. Gondek, Ravindra Majeti, Hongkai Ji, Michael A. Koldobskiy, Andrew P. Feinberg

## Abstract

Disruption of the epigenetic landscape is of particular interest in acute myeloid leukemia (AML) due to its relatively low mutational burden and frequent occurrence of mutations in epigenetic regulators. Here, we applied an information-theoretic analysis of methylation potential energy landscapes, capturing changes in mean methylation level and methylation entropy, to comprehensively analyze DNA methylation stochasticity in subtypes of AML defined by mutually exclusive genetic mutations. We identified AML subtypes with CEBPA double mutation and those with IDH mutations as distinctly high-entropy subtypes, marked by methylation disruption over a convergent set of genes. We found a core program of epigenetic landscape disruption across all AML subtypes, with discordant methylation stochasticity and transcriptional dysregulation converging on functionally important leukemic signatures, suggesting a genotype-independent role of stochastic disruption of the epigenetic landscape in mediating leukemogenesis. We further established a relationship between methylation entropy and gene expression variability, connecting the disruption of the epigenetic landscape to transcription in AML. This approach identified a convergent program of epigenetic dysregulation in leukemia, clarifying the contribution of specific genetic mutations to stochastic disruption of the epigenetic and transcriptional landscapes of AML.

## Main Text

Acute myeloid leukemia (AML) is a hematopoietic malignancy driven by a combination of genetic lesions and epigenetic disruptions, leading to abnormal clonal expansion and blockade of myeloid differentiation (*1, 2*). While AML has a relatively low mutational burden, mutational analysis has identified recurrent genetic lesions in epigenetic machinery like *DNMT3A, IDH1* or *IDH2, TET2*, and *MLL* (*KMT2A*); in transcription factors like *CEBPA* and *RUNX1*; and in signaling molecules such as *FLT3, KIT*, and *NRAS* (*1, 2*). Interestingly, mutations in the epigenetic machinery often occur early in leukemogenesis, suggesting an important role of epigenetic regulation in leukemia (*3, 4*). Previous work has identified specific patterns of DNA methylation linked to mutational profiles in AML, such as hypomethylation in DNMT3A-mutant AMLs and hypermethylation in IDH-mutant AMLs (*5*–*8*). Analysis of DNA methylation data has also revealed that some genetic subtypes of AML may be distinguished by methylation signatures, particularly subtypes with mutations in epigenetic machinery, and that this classification may hold prognostic relevance (*9*–*13*). Thus, the role of mutations in the epigenetic machinery in mediating disruption of the epigenetic landscape of AML is a crucial topic of investigation.

We and others have shown methylation stochasticity to be linked to plasticity, tumor evolution, and prognosis in leukemia and other cancers (*14*–*22*). To rigorously capture the higher-order statistical properties of methylation, our group has previously developed information-theoretic methods to model methylation potential energy landscapes (PELs), enabling the detection of stochastic disruption of the epigenetic landscape (*23*–*26*). These methods allow for direct comparisons of the underlying probability distributions of methylation, measuring differential methylation stochasticity by capturing changes in both mean methylation level and methylation entropy, as well as other properties that are not detectable by mean-based analysis. We have previously applied these information-theoretic methods to investigate epigenetic landscape disruption in solid tumors (*14, 23*), in acute lymphoblastic leukemia (ALL) (*16*), and in MLL-rearranged AML (*15*), where we identified discordant methylation stochasticity over key regulators of the malignant phenotype, pointing to the role of stochastic epigenetic regulation in oncogenesis. Li et al. subsequently carried out a comparative analysis of epigenetic heterogeneity in genetic subtypes of AML using empirical metrics of epiallele diversity, finding that specific somatic driver mutations were associated with epigenetic allele diversity over distinct loci (*22*). To further dissect the contribution of AML driver mutations to disruption of the epigenetic landscape, we applied the Correlated Potential Energy Landscape (CPEL) method (*25, 26*) to characterize the methylation PELs of genetically defined AML subtypes, focusing on subtypes with mutually exclusive mutations in *DNMT3A, IDH1/2, TET2, CEBPA* double mutation (*CEBPA*-dm), *KIT*, or *NRAS*. We sought to carry out a comprehensive analysis of methylation stochasticity in genetically-defined AML subtypes, parsing out the contribution of individual mutational drivers to the disruption of the epigenetic landscape. By integrating epigenetic landscapes with single-cell RNA sequencing, we investigated a functional relationship between DNA methylation stochasticity and gene expression.

### Information-theoretic methylation potential energy landscape analysis reveals discordant methylation stochasticity in AML

To investigate methylation stochasticity driven by genetic mutations in AML, we applied CPEL, an information-theoretic method for analysis of differences in methylation potential energy landscapes (*26*). CPEL detects differences in mean methylation level (MML), normalized methylation entropy (NME), and probability distributions of methylation (PDMs) based on methylation patterns within analysis regions (**Figure 1A, S1**). Differences in PDMs are quantified by the uncertainty coefficient (UC), the geometric Jensen-Shannon divergence normalized by the cross-entropy between two groups, which captures both mean and entropy changes (**Methods**). We evaluated analysis regions containing an average of 6.5-9 CpGs per region (**Data S1**), requiring CpGs to have at least 10x coverage in every sample (**Methods**).

**Figure 1:**
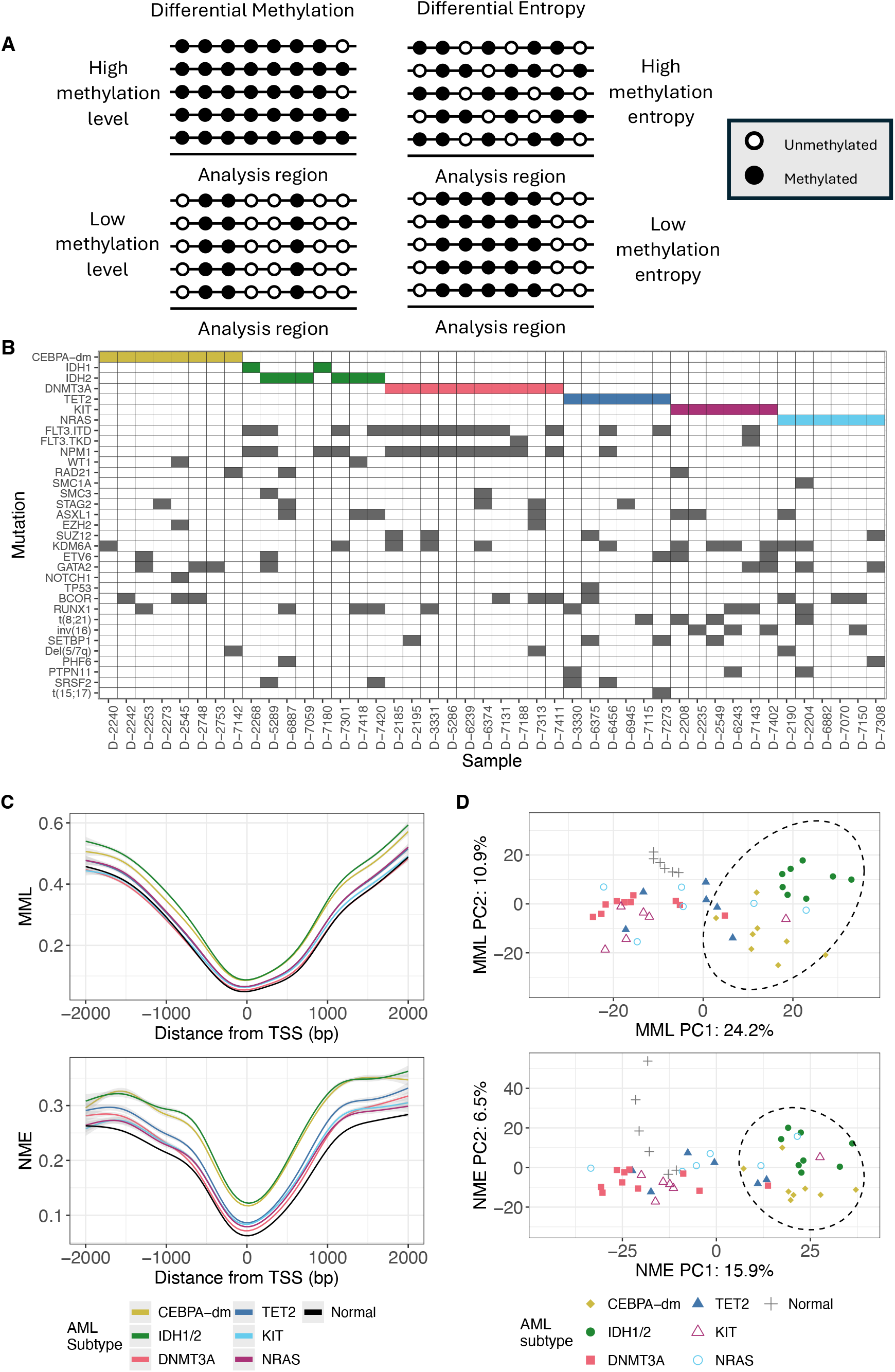
Potential energy landscape analysis identifies discordant methylation stochasticity in AML. **(A)** Illustration of differential mean methylation (left) and differential methylation entropy (right). Both cases have differences in probability distributions of methylation, but one is driven by differences in mean methylation level (left) and the other by differences in methylation entropy (right). **(B)** Mutational profiles (rows) of selected ERRBS samples (columns) used in CPEL analysis. Mutations in genes used for definition of AML subtypes are indicated by color, and are mutually exclusive (CEBPA-dm, yellow; IDH1/2, green; DNMT3A, pink; TET2, dark blue; NRAS, purple; KIT, light blue). **(C)** Meta-region plots of CPEL MML (top) and NME (bottom) over promoter regions, colored by AML subtype. **(D)** Principal component analysis (PCA) of CPEL MML (top) and NME (bottom), colored by AML subtype.

We employed a dataset of enhanced reduced representation bisulfite sequencing (ERRBS) data from a large cohort of primary AML samples (*6*). Samples were selected based on the presence of mutually exclusive mutations in *DNMT3A* (n=10), *IDH1/2* (n=8), *TET2* (n=6), *CEBPA* double mutation (CEBPA-dm, n=8), *KIT* (n=6), or *NRAS* (n=6), allowing for evaluation of epigenetic landscape disruption mediated by specific somatic mutations without the presence of confounding mutations of other subtypes (**Figure 1B, Data S2**). ERRBS data from healthy CD34+ progenitors was used for normal reference (n=6) (*27, 28*). Strikingly, CEBPA-dm and IDH1/2-mut AMLs had distinct hypermethylation and increased methylation entropy over promoter regions (**Figure 1C, S2-3**). Principal component analysis (PCA) of MML and NME separated CEBPA-dm and IDH1/2-mut AMLs from other subtypes and normal samples (**Figure 1D**), indicating that these two subtypes have strong and convergent methylation disruption not found in other AML subtypes. DNMT3A-mut and TET2-mut AMLs had modest methylation stochasticity discordance (3,358 and 2,752 regions with significant UC, respectively) (**Data S1**), despite the canonical role of these enzymes as mediators of DNA methylation (*29, 30*).

We observed strong disruption of methylation over CpG islands (CGI), shores, and shelves (**Figure S2, S3**). Regions with significant differences in PDMs (UC DMRs) were enriched over CGI and promoter regions in CEBPA-dm, IDH1/2, TET2, KIT, and NRAS-mut AMLs (**Figure S4**). Interestingly, UC DMRs in DNMT3A-mut AMLs were depleted over these features and enriched over open seas and repetitive/CNV elements, indicating that DNMT3A mutations may mediate changes in DNA methylation outside of CGI-associated features. UC DMRs were also enriched in heterochromatin regions, suggesting a role of increased methylation stochasticity in disruption of chromatin organization.

HOMER motif analysis of DMRs revealed convergent enrichment of UC DMRs in CEBPA-dm and IDH1/2-mut AMLs over transcription factor binding motifs for Homeobox family transcription factors such as HOXA1, HOXC6, CUX1, and OCT4 (*POU5F1*) (**Figure S5, Data S4**). CEBPA-dm and IDH1/2-mut DMRs also both showed enrichment over motifs for AP-2 (*TFAP2A, TFAP2G*), and LRF (*ZBTB7A*) transcription factors. These transcription factors play important roles in AML: disruption of Homeobox family members in AML is well-established (*31*); AP-2 transcription factors have been implicated in disease progression through *HOXA* disruption (*32*); and *ZBTB7A* is necessary for myeloid differentiation and is frequently mutated in *t(8;21)* AML (*33*). In contrast, DMRs in DNMT3A-mut AMLs were enriched over motifs for ETS, RUNX, and bZIP family transcription factors, such as PU.1 (*SPI1*), a key regulator in hematopoiesis (*34*); RUNX2, implicated in differentiation and development of *Cbfb-MYH11* AML (*35*); and AP-1 (*JUNB*), which has been linked to development of myeloproliferative disease in mice (*36*) (**Figure S5**). Thus, genetic mutations, particularly mutations in epigenetic machinery, may contribute to leukemogenesis by mediating disruption of DNA methylation over binding motifs for key transcription factors.

Together, these results indicate that genetic mutations are associated with highly stochastic methylation in AML. Across subtypes, differential methylation and disruption of transcription factor binding motifs localizes to key regulatory elements involved in leukemia. Furthermore, we find that CEBPA-dm and IDH1/2-mut AMLs have distinct and highly entropic methylation profiles. Both of these mutations have previously been associated with hypermethylation in AML (*2, 9*). Li et al. identified characteristic distribution of epigenetic allele diversity associated with CEBPA-dm AML, but observed less tightly linked profiles in IDH1/2-mut AMLs (*22*). The stochastic methylation landscapes of these subtypes suggests a crucial role of epigenetic regulation and disruption of methylation in AML.

### Discordant methylation stochasticity localizes to key regulators of the leukemic phenotype across AML subtypes

Next, we mapped regions with significant differences in PDM (UC DMRs), MML (dMML DMRs), or NME (dNME DMRs) to genes (**Data S3**). CEBPA-dm and IDH1/2-mut AMLs had strong convergence of genes containing DMRs (**Figure 2A, S6A-B**), indicating that the increased methylation stochasticity observed in these AML subtypes occurs over similar targets. The overlap between UC DMR genes in CEBPA-dm and IDH1/2-mut AMLs consisted of 4,090 genes, including *SETBP1, UHRF1*, and *MEIS2* (**Data S5**), all of which play roles in epigenetic regulation, hematopoiesis, or leukemia development (*37*–*39*). Our group has previously shown convergent methylation discordance over *UHRF1* across cytogenetic subtypes of ALL (*16*). We also identified a set of 24 genes with UC DMRs in all AML subtypes (**Data S5**). This gene set included *IRX2*, a member of the Iroquois homeobox gene family previously found to be predictive of outcome in infant ALL (*40*); *DOK6*, for which promoter methylation may serve as a prognostic biomarker in AML (*41*); and *DUSP1*, which is linked to therapy resistance in *BCR-ABL1* chronic myeloid leukemia (*42*).

**Figure 2:**
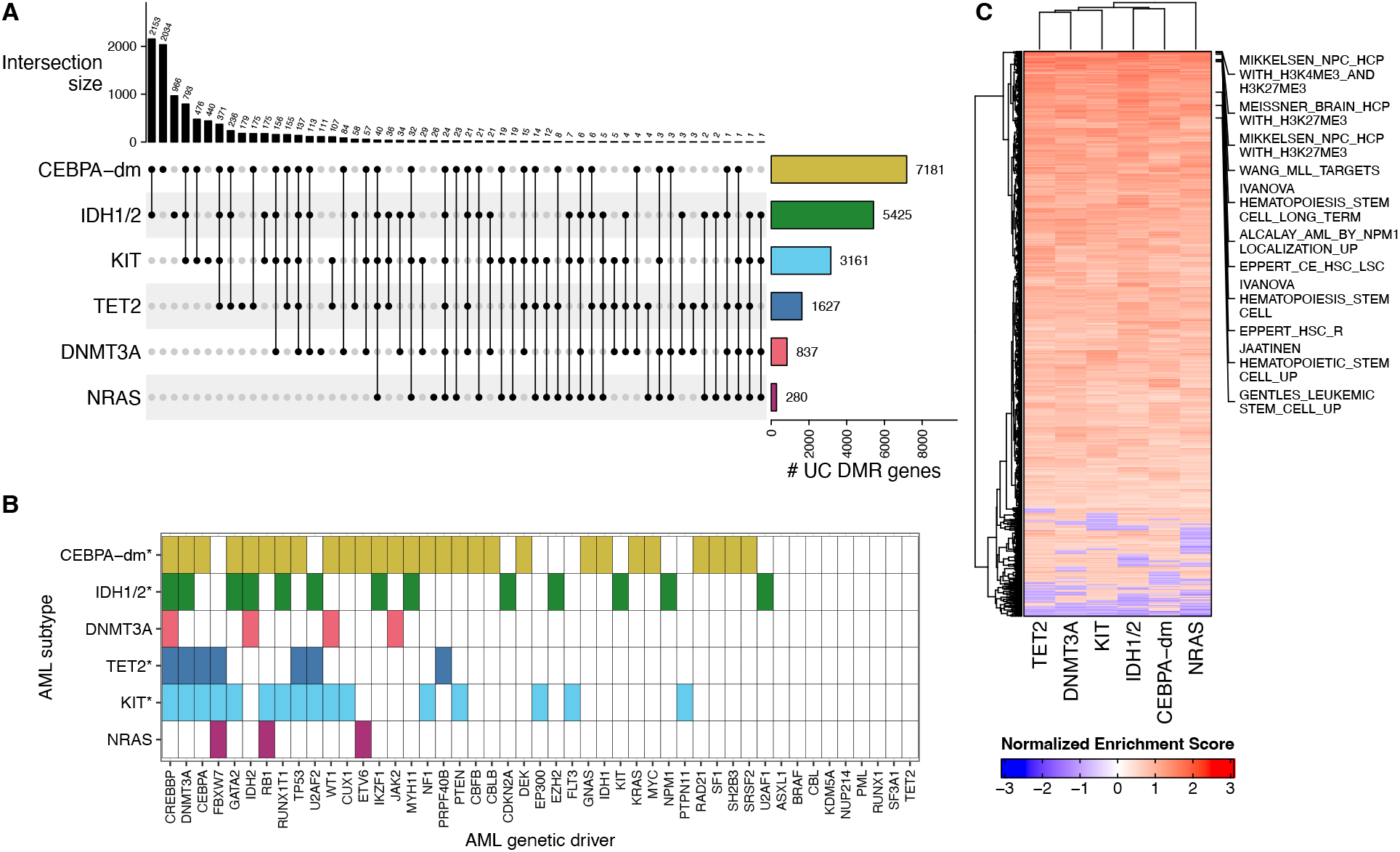
Methylation discordance in AML localizes to key regulators of leukemia. **(A)** Overlap of genes associated with UC DMRs between AML subtypes. Genes with associated analysis regions are listed in **Data S3. (B)** Genetic drivers of AML have disruption of methylation (measured by CPEL UC) in AMLs, independent of mutations in that genetic driver. Colors indicate the AML subtype (rows) with methylation disruption over the promoter of the given gene (columns). AML genetic drivers covered by CPEL analysis regions are included. *: significant enrichment of AML genetic driver genes in UC DMR genes for the given subtype (hypergeometric p-value < 0.01). **(C)** Gene set enrichment analysis of genes ranked by promoter UC over the MSigDB C2 gene set library (**Methods**). Select gene sets relevant to histone marks and leukemic signatures are annotated. Positive NES = high UC (highly discordant methylation stochasticity); negative NES = low UC (lowly discordant methylation stochasticity).

Surprisingly, DNMT3A-mutant AMLs had fewer genes associated with DMRs than TET2-mutants (837 versus 1,627 UC DMR genes; **Figure 2A**), despite having a greater number of DMRs in all (3,358 versus 2,752 UC DMRs; **Data S1**). This suggests that methylation discordance mediated by *DNMT3A* mutations in AML occurs mostly in non-promoter regions, a result that is supported by the unique enrichment of DNMT3A DMRs in open seas and repetitive elements (**Figure S4**). Previous reports have also found that methylation changes mediated by *DNMT3A* mutations target non-CGI features (*43*), corroborating these results.

We observed that the set of genes associated with UC DMRs in CEBPA-dm and IDH1/2-mut AMLs contained genes recurrently mutated in AML. Based on this observation, we explored whether other subtypes also exhibited methylation discordance over genes commonly altered in AML. We generated a list of genetic drivers of AML, encompassing genes frequently mutated involved in translocations in AML (*1*); 47 genetic drivers were covered in the CPEL analysis. Surprisingly, all AML subtypes demonstrated increased methylation stochasticity over key drivers of AML, even in the absence of mutations in those genes (**Figure 2B**). In total, 38 of the 47 genetic drivers had increased methylation stochasticity in at least one AML subtype. We found significant enrichment of AML driver genes in the list of genes with UC DMRs for CEBPA-dm, IDH1/2-mut, KIT-mut, and TET2-mut AMLs (hypergeometric p < 0.01). This suggests that drivers of AML may be disrupted either genetically or epigenetically, thus contributing to leukemogenesis. This result mirrors previous work in ALL, which revealed significant methylation discordance over chromosomal translocation genes across ALL cases, independent of cytogenetic status (*16*).

Next, we performed over-representation analysis (ORA) and gene set enrichment analysis (GSEA) of genes with DMRs. ORA revealed enrichment of genes with UC DMRs in targets of PRC2 complex subunits (*EZH2, SUZ12*) and other transcription factors such as *REST, TRIM28*, and *FOXA1*, which are involved in pluripotency and chromatin organization (*44*–*48*). Genes with UC DMRs in CEBPA-dm and IDH1/2-mut AMLs were also strongly enriched in *NANOG* targets, and all subtypes except DNMT3A- and NRAS-mut AMLs were enriched in *SMAD4* targets (**Figure S6C, Data S6**). GSEA of the Molecular Signatures database (MSigDB) C2 collection revealed that genes with high UC were enriched in gene sets involving hematopoietic stem cell function, leukemia signatures, and H3K27me3 or H3K4me3 marks (**Figure 2C, Data S7**). We observed enrichment in MSigDB Hallmark gene sets relevant to oncogenesis such as epithelial-to-mesenchymal transition, hypoxia, IL2-STAT5 signaling, and KRAS signaling (**Figure S6E, Data S8**). Genes with dMML or dNME DMRs showed similar enrichment in targets of the PRC2 complex and transcription factors regulating pluripotency, as well as in gene sets relevant to leukemogenesis and chromatin marks (**Figure S6C-E, Data S6-8**).

These results demonstrate convergent disruption of methylation over crucial drivers of leukemia across all AML subtypes, regardless of genetic mutation. The enrichment of genes with high methylation stochasticity in gene sets relevant to leukemia biology suggests that stochastic epigenetic landscape disruption occurs over a core set of leukemia regulators, potentially driving leukemogenesis across AMLs. CEBPA-dm and IDH1/2-mut AMLs had distinctive profiles of methylation stochasticity, indicating that these subtypes have additional factors mediating epigenetic disruption. Together, these findings highlight the role of methylation stochasticity in shaping the epigenetic landscape to mediate leukemic transformation.

### Transcriptional dysregulation in AML occurs over epigenetically-disrupted regulators of leukemia

To assess the influence of DNA methylation on gene expression in AML, we performed single-cell RNA sequencing on a cohort of primary AMLs obtained from Johns Hopkins Hospital and Stanford University (**Methods**). This cohort included AMLs with mutations in *DNMT3A* (n=5), *IDH1/2* (n=4), *TET2* (n=7), *CEBPA*-dm (n=5), plus healthy CD34+ progenitors (CD34+ and GMP) for normal reference (n=3) (**Data S2**). Uniform manifold approximation and projection (UMAP) analysis of scRNA-seq data identified ten clusters. Samples were dominant in either Cluster 0 (progenitor-like) or Cluster 1 (monocyte-like) (**Figure 3A, S7**). All CEBPA-dm samples were Cluster 0-dominant, suggesting that these AMLs have an undifferentiated, progenitor-like expression profile (**Figure S7B-C**).

**Figure 3:**
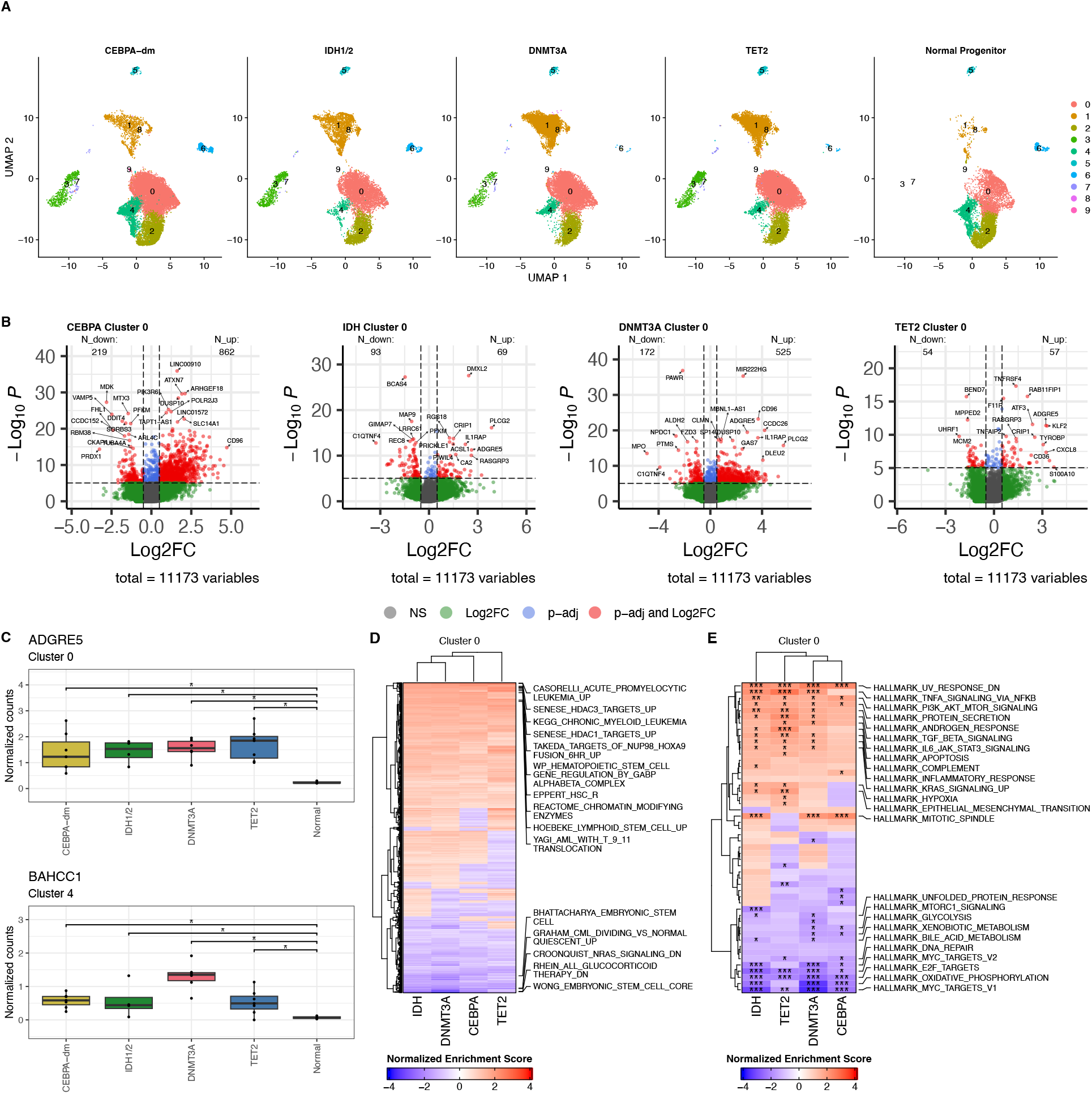
Single-cell RNA-seq reveals transcriptional dysregulation over leukemia signatures in AML. **(A)** UMAPs of scRNA-seq revealing 10 distinct clusters. Samples are pooled by genotype. **(B)** Volcano plots of differentially expressed genes in Cluster 0 for CEBPA-dm AMLs (left), IDH1/2-mutant AMLs (middle-left), DNMT3A-mutant AMLs (middle-right), and TET2-mutant AMLs (right) versus normal progenitors. **(C)** Boxplots of cluster-level pseudobulk expression profiles for *ADGRE5* (in cluster 0) and *BAHCC1* (in cluster 4), differentially expressed in all AML versus normal comparisons. *: Bonferroni adjusted p-value ≤ 0.05. **(D)** Gene set enrichment analysis over the MSigDB C2 gene set library of genes ranked by average log2 fold change (versus normal samples) in cluster 0. Selected gene sets related to histone marks, leukemia, and hematopoietic signatures are annotated. **(E)** Gene set enrichment analysis over the MSigDB Curated Hallmark gene set library of genes ranked by average log2 fold change (versus normal samples) in cluster 0. *: Benjamini-Hochberg adjusted p-value ≤ 0.05; **: adjusted p-value ≤ 1e-3; ***: adjusted p-value ≤ 1e-4. Positive NES = upregulated in AML relative to normal samples; negative NES = downregulated in AML relative to normal.

First, we performed differential expression analysis between each AML subtype and normal progenitors within UMAP clusters 0, 2, and 4 (**Methods**). These clusters are of particular interest due to their progenitor-like expression (**Figure S7A**). We observed the strongest transcriptional dysregulation in CEBPA-dm AMLs (1,081; 863; and 652 differentially expressed genes (DEGs) in clusters 0, 2, and 4 respectively) (**Figure 3B, S8A-B**; **Data S9**). Surprisingly, despite the high methylation stochasticity observed in IDH-mutant AMLs, this subtype had minimal transcriptional disruption (162; 178; and 33 DEGs in clusters 0, 2, and 4 respectively), while DNMT3A-mutant AMLs, despite having only modest changes in methylation landscape, had stronger dysregulation of gene expression (697; 401; and 614 DEGs in clusters 0, 2, and 4 respectively) (**Figure 3B, S8A-B**; **Data S9**).

We identified five genes with differential expression across all AML subtypes, including *ADGRE5* (*CD97*) upregulated in cluster 0 and *BAHCC1* upregulated in cluster 4 (**Figure 3C, S8C**). Most of these genes had discordant methylation stochasticity in AMLs; for example, *BAHCC1* was associated with UC DMRs in CEBPA-dm, TET2-mutant, and KIT-mutant AMLs (**Data S3**). In fact, 4 of the 5 common differentially expressed genes had DMRs in CEBPA-dm AMLs. *ADGRE5* has been identified as a crucial regulator of leukemic stem cells (LSCs) (*49*), and *BAHCC1* was reported to sustain leukemogenesis through its function as a reader of the Polycomb mark H3K27me3 (*50*). Known genetic drivers of AML were also differentially expressed in CEBPA-dm and DNMT3A-mutant AMLs (**Data S10**). For example, *CBLB, JAK2* and *KDM6A* were upregulated in both CEBPA-dm and DNMT3A-mutant AMLs in clusters 0, 2, and 4 (**Figure S8D**). Several of these genes had increased methylation stochasticity, such as *JAK2* in CEBPA-dm and DNMT3A-mutant AMLs (**Figure 2B**).

ORA revealed convergent enrichment of DEGs in targets of the PRC2 complex (*SUZ12*) and in *SMAD4* targets across clusters 0, 2, and 4 (**Data S11**). This parallels the enrichment of PRC2 and *SMAD4* targets observed in DMR associated genes (**Figure S6C**). In agreement with the enrichment over genes with high methylation stochasticity (**Figure 2C, S6D-E**), GSEA revealed convergent enrichment of DEGs in all AML subtypes over leukemia signatures from the MSigDB C2 collection (**Figure 3D, S8E**; **Data S12**). We also observed a consistent upregulation of gene sets relevant to oncogenesis from the MSigDB Hallmark collection, including PI3K/AKT, TGF-beta, and KRAS signaling (**Figure 3E, S8F**; **Data S13**). Interestingly, we MYC gene sets from the MSigDB Hallmark collection were downregulated across AML subtypes in all clusters analyzed, despite the known role of MYC deregulation in leukemogenesis (*51*).

Taken together, these results demonstrate the transcriptional disruption of key leukemia regulators across AML subtypes. The convergence of discordant methylation stochasticity and transcriptional dysregulation over similar gene targets and leukemia signatures highlights the important role of epigenetic landscape disruption on the transcriptional phenotype in AML, suggesting that transcriptional changes linked to methylation stochasticity may be a defining feature of leukemogenesis.

### DNA methylation stochasticity is associated with transcriptional dysregulation in AML

To further investigate the connection between methylation discordance and transcriptional disruption in AML, we computed the mean-adjusted variability (MAV), a measure of stochastic gene expression, for each detected gene in each sample (**Methods, Data S14**). We observed a strong relationship between NME near the TSS and quartiles of genes ranked by MAV, where genes with higher MAV had increased NME near the TSS (**Figure 4A, S9**). Gene expression mean was inversely related to mean methylation near the TSS, where genes with higher mean expression had lower MML near the TSS. This result links DNA methylation stochasticity to transcriptional variability, providing an explanation for the convergent enrichment of the two modes over key regulators of leukemia, and suggesting how these changes in DNA methylation may affect the transcriptional and therefore phenotypic state of AML.

**Figure 4:**
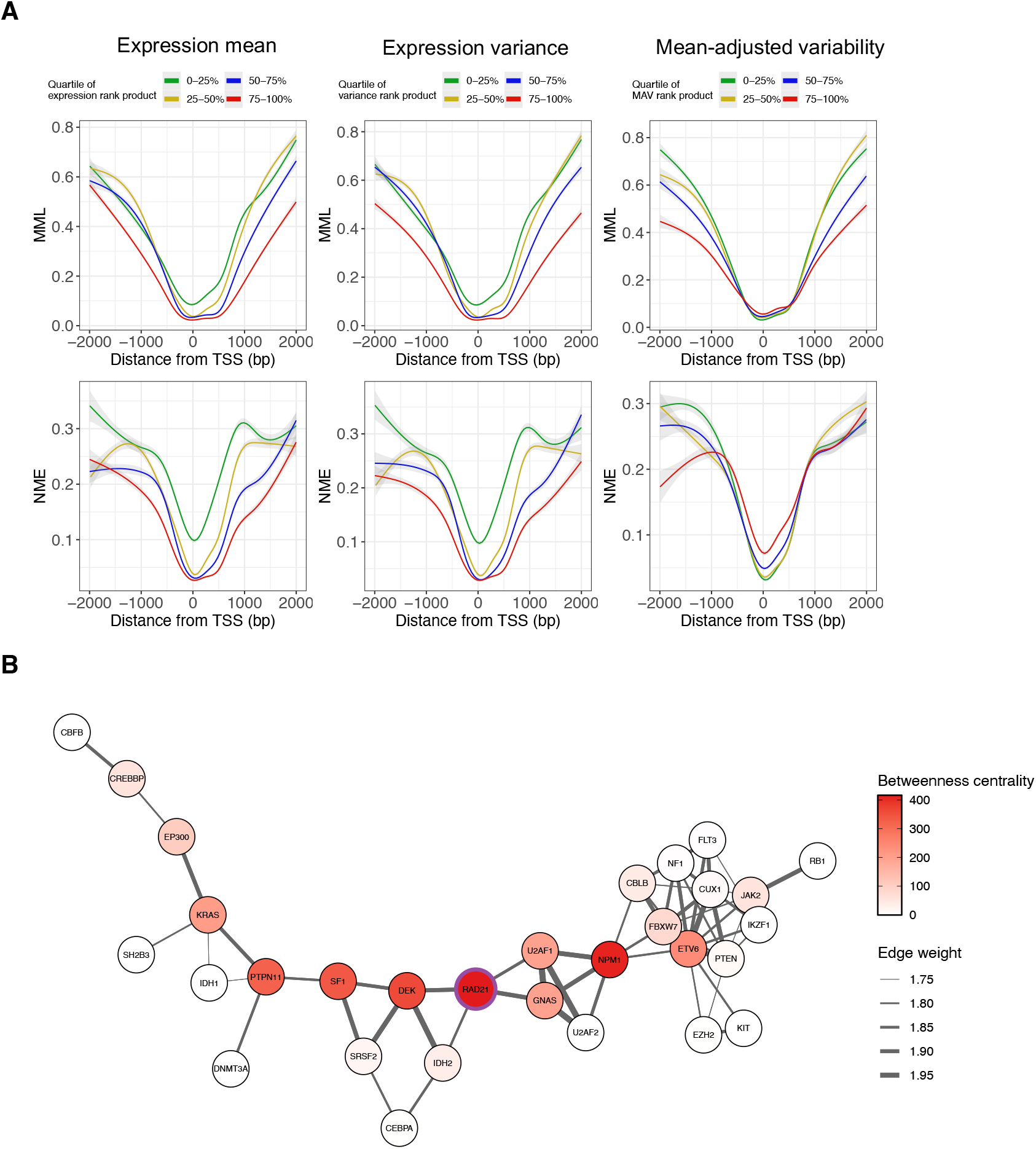
DNA methylation stochasticity is associated with gene expression in AML. **(A)** Relationship between MML or NME and gene expression mean (left), variance (middle), or variability (MAV; right) in IDH1/2-mut AMLs as a function of distance from the TSS. Lower methylation level is associated with higher expression level (top-left), while lower expression levels are associated with lower levels of NME (bottom-left). Mean-adjusted expression variability is associated with higher NME near the TSS, connecting methylation stochasticity to expression variability. **(B)** Information-theoretic analysis of scRNA-seq data reveals a complex gene regulatory network between 30 genetic drivers of AML. All included genetic drivers show disrupted methylation landscapes by the CPEL model. The gene with the highest betweenness centrality (*RAD21*) is marked.

Finally, we employed a multivariate information-theoretic method of gene regulatory network inference (*52*) to explore dependencies among the 38 genetic drivers of AML with significantly discordant methylation stochasticity (**Figure 2B**). We assessed CEBPA-dm, IDH1/2-mut, and TET2-mut AMLs, as these subtypes showed statistically significant overlaps between genetic drivers of AML and genes with significant methylation stochasticity. This analysis revealed a complex regulatory network among 30 of these drivers of AML (**Figure 4B**). In this network, *RAD21* was predicted to have the highest betweenness centrality, indicating that it exerts the greatest influence over the network. Interestingly, *RAD21* was directly connected to *IDH2*, and *IDH2* was directly connected to *CEBPA*, the genes defining AML subtypes with markedly high methylation stochasticity. *RAD21* is a subunit of the cohesin complex, with an essential role in mediating chromatin structure through formation of loops and topologically-associating domains (TADs) (*53*). The central position of this chromatin regulator in the gene regulatory network suggests that epigenetic and transcriptional dysregulation of key mediators of genome organization may influence leukemogenesis.

These results show that high methylation stochasticity is linked to transcriptional dysregulation in AML, with increased methylation entropy associated with higher gene expression variability. Additionally, we identify a gene regulatory network between non-mutated AML driver genes containing DMRs, suggesting that key drivers of AML may be genetically or epigenetically disrupted in leukemia. Together, these findings provide evidence of how methylation discordance may directly contribute to transcriptional profiles driving leukemogenesis.

## Discussion

Methylation stochasticity has been shown to be associated with clinical outcome, chemoresistance, and disease progression in hematopoietic malignancies (*16, 18*–*22*). A recent study reported that somatic mutations are associated with epigenetic heterogeneity in AML, suggesting a relationship between increased epigenetic variability and inferior prognosis (*22*). Surprisingly, in this study IDH1/2 mutant AML had broad distributions of epialleles, without characteristic clustering as seen for other genetic subtypes of AML (*22*). To investigate the contribution of specific, mutually-exclusive genetic mutations to stochastic disruption of the epigenetic landscape, we conducted an information-theoretic analysis of methylation potential energy landscapes in genetic subtypes of AML. Our analyses identified a core program of methylation stochasticity and transcriptional disruption involving key regulators of leukemia biology across all AMLs. Furthermore, we link methylation entropy to gene expression variability, suggesting that transcriptional dysregulation associated with methylation stochasticity may be a hallmark of leukemogenesis, regardless of genetic mutations. All subtypes of AML had significant methylation discordance over genetic drivers of AML, even in the absence of mutations in those genetic drivers. These genes formed a complex gene regulatory network, indicating that disruption of genetic drivers of AML by methylation stochasticity may drive dysregulation of the transcriptional state of crucial regulators of leukemia. We additionally found that CEBPA-dm and IDH1/2-mutant AMLs have markedly increased DNA methylation entropy beyond this core leukemia program, suggesting that these subtypes have additional factors driving dysregulation of the epigenetic landscape.

Our observation of markedly increased methylation entropy in CEBPA-dm and IDH1/2-mutant AMLs is in agreement with findings by Li et al. (*22*), who reported a high degree of epigenetic heterogeneity in specific AML subtypes including CEBPA-dm and IDH1/2-mutated. Here, we describe a distinct and surprisingly convergent profile of methylation stochasticity in both CEBPA-dm and IDH1/2-mut AMLs. This complements previous findings of increased epiallele heterogeneity in these AML subtypes by providing gene-level resolution, demonstrating the sensitivity of information-theoretic methods for methylation PEL analysis, which can capture higher-order statistical properties of methylation that may not be detected by empirical estimations of methylation entropy (*23*–*25*). In CEBPA-dm and IDH1/2-mut AMLs, we observed convergent enrichment of Homeobox, AP-2, and LRF transcription factor binding motifs over regions with stochastic methylation disruption. These transcription factors are known to mediate leukemogenesis: overexpression or translocations involving Homeobox genes contribute to malignancy (*31*); AP-2 can negatively regulate *CEBPA* and may contribute to *HOX* dysregulation in AML (*32, 54*); and LRF (*ZBTB7A*) is a coordinator of hematopoietic differentiation (*33*). At the gene level, methylation landscape disruption in CEBPA-dm and IDH1/2-mutant AMLs was also highly convergent, occurring over genes such as *SETBP1, UHRF1*, and *MEIS2*. These genes play crucial roles in epigenetic regulation, leukemia, or hematopoiesis: *SETBP1* recruits chromatin modifiers to control gene expression, and is frequently mutated in leukemia (*37*); *UHRF1* coordinates DNA methyltransferase *DNMT1* to repressive chromatin marks, and was identified as a convergent target of stochastic DNA methylation in ALL (*16, 38*); and *MEIS2* is an important factor in myeloid differentiation (*39*).

CEBPA- and IDH-mutant AMLs have previously been characterized as hypermethylated; *IDH* mutations in particular lead to a well-described hypermethylated profile (*2, 7, 8*). Mutant *IDH* enzymes produce the oncometabolite 2-hydroxyglutarate, which competitively inhibits alpha-ketoglutarate-dependent enzymes, including *TET* family members and histone lysine demethylases (*55*). The effect of *IDH* mutations on prognosis of AMLs is unclear, and may depend on co-occurring mutations, treatment, and other patient characteristics (*30*). Additionally, small-molecule inhibitors of *IDH1* or *IDH2* may be used in the treatment of IDH-mutant AMLs; these been reported to reduce methylation levels in human AML (*55*) and to lead to decreased epigenetic allele diversity in a mouse model of AML (*22*). AMLs with biallelic or bZIP-domain mutations in *CEBPA* are a distinct subtype classified by the World Health Organization (*56*), and are associated with a favorable prognosis (*57*). Interestingly, mutations in *CEBPA* have been associated with resistance to IDH inhibitors (*55*). Aside from its canonical function as a transcription factor, *CEBPA* is also known to interact with epigenetic machinery, such as the histone acetyltransferase p300 and the SWI/SNF complex (*58*–*60*). Recent work has also implicated CEBPA in the control of DNA methylation through direct interaction with DNMT3A, where the interaction between wild-type CEBPA and DNMT3A blocks association with DNA; this interaction was shown to be lost upon *CEBPA* mutation, leading to aberrant hypermethylation (*61*). The disruption of interactions between *CEBPA* and epigenetic regulators upon *CEBPA* mutation may provide a basis for the observed increase in methylation stochasticity, potentially contributing to the highly disordered epigenetic landscape of CEBPA-dm AMLs. A similar process may underlie the defined patterns of epigenetic alleles previously observed in AMLs with other canonical transcription factor mutations, such as t(8;21), inv(16), and t(15;17) (*22*). The disparate prognostic impact of *CEBPA* and *IDH1/2* mutations indicates that increased methylation stochasticity is not always a marker of unfavorable outcome; rather, it may reflect widespread deregulation of the epigenetic machinery, whether by oncometabolites (as in the case of *IDH* mutations) or by disruption of crucial protein interactions (as in the case of *CEBPA* mutations).

DNMT3A-mut and TET2-mut AMLs had only modest discordance of methylation stochasticity, despite the well-defined functions of these enzymes as direct modifiers of DNA methylation (*29, 30*). Nevertheless, methylation discordance in these subtypes localized to important regulators of malignancy. It is possible that mutations in these genes mediates changes in DNA methylation outside of regions covered by the ERRBS data analyzed here. Indeed, the enrichment of DMRs in DNMT3A-mut AMLs over open seas suggests that these mutations may predominantly affect non-CGI regions. This observation aligns with previous studies showing differential methylation over open seas (*43*) and an absence of CpG island disruption (*5*) in DNMT3A-mutant AMLs. Likewise, *TET2* mutations have been shown to affect primarily non-CGI methylation (*62*), particularly methylation over enhancer regions (*63*), which may not be captured by ERRBS. Further investigation into the effect of mutations in these enzymes on DNA methylation stochasticity is warranted.

We identified a surprising convergence of discordant methylation stochasticity over key regulators of leukemia and pluripotency across all AML subtypes. These included targets of the PRC2 complex, *REST*, and *TRIM28*, which are factors crucial for epigenetic regulation and pluripotency (*45*–*48*). Genes with methylation discordance were also enriched in gene sets relevant to oncogenesis and leukemia across AML subtypes. Notably, we also observed significantly increased methylation stochasticity over genetic drivers of AML such as *CREBBP, GATA2*, and *RUNX1T1* (*1*) in all AML subtypes, independent of mutations in those driver genes. In contrast, Li et al. found epigenetic heterogeneity over mediators of leukemogenesis only in specific subtypes of AML with particularly high epigenetic allele diversity (*22*). Information-theoretic methods therefore may allow more sensitive detection of stochastic dysregulation of the epigenetic landscape. These results suggest disruption of the epigenetic landscape over critical drivers of leukemogenesis may be a core feature of leukemogenesis across AMLs.

Like methylation stochasticity, transcriptional dysregulation was convergently enriched over key regulators of leukemia across AML subtypes, suggesting a link between methylation disruption and transcriptional dysregulation. We further support this result by demonstrating a relationship between methylation stochasticity and gene expression, where increased methylation levels correlate with lower mean gene expression, and higher methylation entropy is associated with greater gene expression variability. This corroborates the previous observation that higher epiallele diversity corresponded with higher transcriptional variance across AML samples (*22*). The connection between methylation stochasticity and gene expression suggests that methylation discordance may set the stage for aberrant transcriptional programs in leukemia.

We identified a complex gene regulatory network between known genetic drivers of AML. These genes also had stochastic disruption of the methylation landscape, suggesting that epigenetic regulation of genetic drivers of AML, even in the absence of mutations in these genes, may contribute to leukemogenesis. This network was centered on *RAD21*, a subunit of the cohesin complex that is crucial for formation of chromatin loops and TADs (*53*), and is mutated in 3% of AMLs (*1*). In this network, *RAD21* was closely connected to *IDH2* and *CEBPA*, suggesting a role of mutations in these genes in transcriptional disruption of *RAD21*. CEBPA stabilizes binding of RAD21 to maintain looping between enhancers and promoters in breast cancer cells (*64*). The central position of this chromatin regulator in the gene regulatory network suggests that chromatin organization may be disrupted by epigenetic and transcriptional dysregulation of key mediators of genome organization in AML.

In conclusion, we leverage high-coverage ERRBS and single-cell RNA-seq data to analyze DNA methylation stochasticity in distinct genetic subtypes of AML, employing an information-theoretic method to construct potential energy landscapes that encompass the higher-order statistical properties of methylation, including both mean methylation and methylation entropy. This approach identified CEBPA-dm and IDH1/2-mutant AMLs as markedly high-entropy subtypes with distinctive and convergent profiles of epigenetic stochasticity. We further identified a core program of methylation stochasticity and gene expression dysregulation over key regulators of the leukemic phenotype across AML subtypes, along with a relationship between methylation entropy and gene expression variability, suggesting that leukemogenesis may be mediated by stochastic disruption of the epigenetic landscape independent of genotype. Taken together, our results indicate that information-theoretic analysis of DNA methylation can elucidate the role of stochastic epigenetic regulation and provide insights into epigenetic drivers of genetically-defined subtypes of AML.

## Acknowledgements

We thank all patients providing AML specimens for this study. We thank all members of the Majeti, Ji, and Feinberg labs for their helpful input and discussion.

## Author contributions

Conceptualization: EH, WZ, EMM, HJ, MK, APF Methodology: EH, WZ, EMM, LPG, RM, HJ, MK, APF Investigation: EH, WZ, EMM, AI, RT, LPG, RM Visualization: EH, WZ Funding acquisition: APF Project administration: MK, APF Supervision: HJ, MK, APF Writing – original draft: EH, MK, WZ, APF Writing – review & editing: EH, MK, WZ, APF

## Data and materials availability

The ERRBS data used in this paper are publicly available (GSE98350). Single-cell sequencing data will be available in GEO upon publication. Processed data are included as supplementary data files, and other materials required for re-analysis are available upon request.

## Supplementary Materials

Figs. S1 to S9

Data S1 to S14

## Materials and Methods

### ERRBS data download and processing

FASTQ files containing enhanced reduced representation bisulfite sequencing (ERRBS) data from 44 primary patient AMLs were downloaded from GSE98350 (*6*). Patients were annotated by 6 genetically defined AML subtypes, requiring mutually-exclusive mutations in each gene: DNMT3A (n=10), IDH1/2 (n=8), TET2 (n=6), CEBPA double mutation (CEBPA-dm, n=8), KIT (n=6), or NRAS (n=6) (**Data S2**). ERRBS data from 6 healthy CD34+ NBM samples was also downloaded for normal reference (GSE52945, GSE108247) (*27, 28*). Reads were trimmed and quality control was performed using TrimGalore v.0.6.6 (https://github.com/FelixKrueger/TrimGalore, RRID:SCR_011847) with default parameters. Alignment to the hg19 reference genome was performed using Bismark v.0.23.0 (RRID:SCR_005604) (*65*) with default parameters. The resulting BAM files were processed with samtools v.1.18 (RRID:SCR_002105) (*66*) for sorting, deduplication, and indexing. Finally, methylation calls were extracted and CpG_report files were generated using Bismark’s methylation_extractor.

### Definition of analysis regions and potential energy landscape estimation

Methylation analysis was performed using the correlated potential energy landscape (CPEL) model for targeted differential methylation using the Julia package CpelTdm.jl (*26*). CpelTdm requires user-defined analysis regions as an input. These were generated by identifying autosomal regions containing at least 3 consecutive CpGs all passing a given coverage filter; coverage filters implemented were: requiring at least 10x coverage in every sample (all AML samples and normal); requiring at least 10x coverage in at least 4 samples per subtype (each AML subtype and normal); requiring at least 5x coverage in every sample (all AML samples and normal); requiring at least 5x coverage in at least 4 samples per subtype (each AML subtype and normal). Regions over 200 bp were split into *max(1, floor(region width/100))* approximately equally-sized subregions using the subdivideGRanges() function from the Bioconductor package exomeCopy (RRID:SCR_001276). The CPEL model was applied to ERRBS data for each AML subtype versus normal comparison using each coverage filter, run over all resulting regions or regions subset to regions within promoters. The coverage filter requiring at least 10x coverage in every sample with all analysis regions as input was selected for downstream analysis, as this set of CPEL comparisons detected the most significant DMRs in TET2-mut AMLs, which had relatively few DMRs under most coverage filters (0-2752 UC DMRs at q ≤ 0.2) (**Data S1**).

The CPEL model estimates the PDM within an analysis region *R* containing *N* CpG sites as:

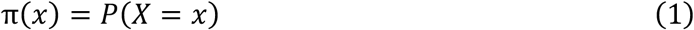

for a random vector ***X***= [*X*_1_ *X*_2_ ‥*X*_*N*_]^*T*^, where X = 0 represents an unmethylated CpG and X = 1 represents a methylated CpG.

After parameter estimation by maximum likelihood method, the mean methylation level (MML) can be computed as:

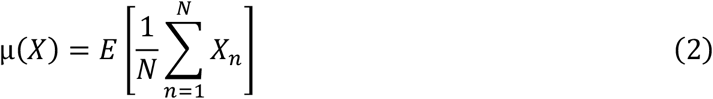

And the normalized methylation entropy (NME) can be computed as:

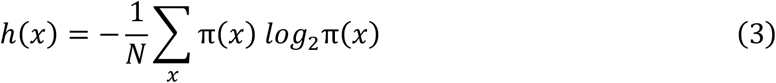

Both MML and NME are normalized to lie within [0, 1], with the highest methylation level taking MML=1 (fully methylated) and the lowest methylation taking MML = 0 (fully unmethylated). Similarly, an NME = 0 indicates there was only one methylation pattern found in the analysis region (fully ordered), and NME = 1 indicates all methylation patterns are equally likely (fully disordered).

### Differential methylation, methylation entropy, and probability distribution of methylation

For a group with *M*_*1*_ and another group with *M*_*2*_ values of CPEL statistics over an analysis region *R*, the CPEL model calculates differential mean methylation level (dMML) with the test statistic:

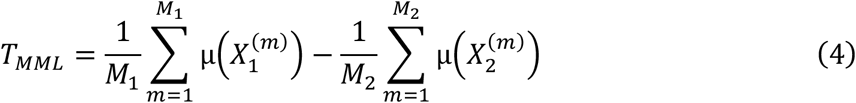

Differential normalized methylation entropy (dNME) is calculated using the test statistic:

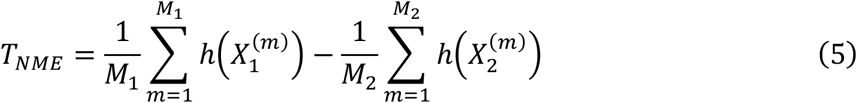

Differences between PDMs are quantified using the uncertainty coefficient (UC) with the test statistic:

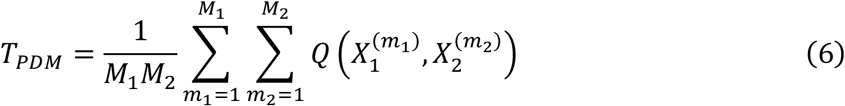

where 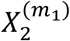 and 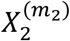 are the m-th methylation states in the first and second group over the analysis region, respectively. Here, the uncertainty coefficient is the geometric Jensen-Shannon divergence normalized by the cross-entropy between groups, given by:

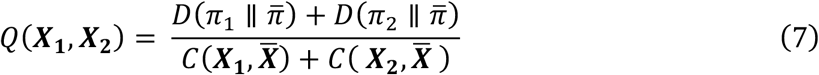

where 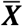 is a random vector associated with the potential energy function 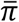, the average of the potential energy functions associated with ***X***_1_ and ***X***_2_; the Kullback-Leibler divergence between two distributions *f* and *g* is

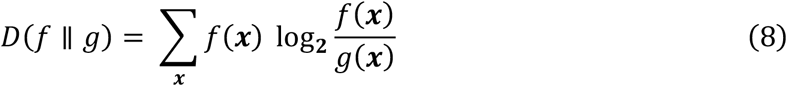

and the cross-entropy between two random vectors ***X***and ***Y*** associated with *f* and *g* is

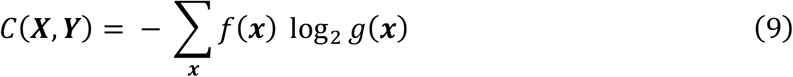

Significance of differential statistics for a given analysis region is determined through permutation-based exact p-value computation or Monte Carlo-based permutation testing (*26*), and p-values are corrected using the Benjamini-Hochberg (BH) procedure for controlling false discovery rate. For downstream analysis, differentially methylated regions (DMRs) were defined as regions with absolute dMML, dNME, or UC ≥ 0.1, and BH-adjusted p-value ≤ 0.2.

### Genomic annotations

Files and tracks have coordinates for hg19. Annotations for CpG islands (CGIs) were obtained from Wu et al. (*67*); CGI shores were defined as 2 kb regions flanking CGIs, CGI shelves were defined as 2 kb regions flanking shores, and open seas were defined as all other genomic regions. Annotations for promoters and gene bodies were obtained from the Bioconductor package TxDb.Hsapiens.UCSC.hg19.knownGene; promoters were defined as ± 2 kb flanking the transcriptional start site. Annotations for K562 chromatin states from ChromHMM (RRID:SCR_018141) and genic features were generated using the R package annotatr’s built-in hg19_K562-chromatin and hg19_basicgenes (DOI: 10.18129/B9.bioc.annotatr).

### Enrichment analysis

For enrichment analysis of DMRs over various genomic features, the odds ratio statistic and Fisher’s two-sided exact test were implemented on a 2 x 2 contingency table containing the number of regions overlapping and not overlapping the feature (such as DMR and overlapping promoter, not DMR and overlapping promoter, etc).

Motif analysis was performed using HOMER v.5.1 (RRID:SCR_010881) with size set to default and CpG normalization method. Input regions were CPEL dMML, dNME, or UC DMRs, and custom background region sets were defined as all CPEL analysis regions.

Over-representation analysis (ORA) and gene set enrichment analysis (GSEA) were implemented using clusterProfiler v.4.12.0 (RRID:SCR_016884). To assess enrichment of genes associated with DMRs or differentially expressed genes over transcription factor target sets, ORA was performed using the gene set library “ENCODE_and_ChEA_Consensus_TFs_from_ChIP-X” downloaded from Enrichr (https://maayanlab.cloud/Enrichr/, RRID:SCR_001575). For DMRs, a background gene set was defined as any gene containing a CPEL analysis region within its promoter; for DEGs, a background gene set was defined as all genes detected in the RNA-seq data. GSEA was performed by ranking genes by the strength of the UC, dNME, or dMML of analysis regions within the gene promoter; or by log2FC reported by Seurat DEseq2 differential expression analysis (see <u>Differential expression analysis</u>). When ranking by CPEL statistics (UC, dNME, dMML), most genes contained multiple analysis regions in the promoter; in this case, the region with the strongest value was used for ranking. GSEA was performed over the MSigDB Hallmark (H) and Curated (C2) gene set collections (Broad Institute, UCSD).

### Single-cell sequencing

Primary patient AML samples for single-cell sequencing experiments were obtained from Johns Hopkins Hospital and Stanford University biobank. Two healthy CD34+ samples were purchased from AllCells (Lot 3011884, 3011592) and one GMP sample was obtained from Stanford University for normal comparison. Samples were collected from peripheral blood or bone marrow; blast percentages ranged from 70-99%. Patients were annotated by 4 genetically defined AML subtypes with mutually exclusive mutations, corresponding to those used for methylation analysis: DNMT3A (n=5), IDH1/2 (n=4), TET2 (n=7), CEBPA-dm (n=5) (**Data S2**).

Cells were thawed in warmed RPMI + 10% FBS, then washed twice with 1X PBS + 0.04% BSA and strained using a 40 μm Flowmi Cell Strainer (SP Bel-Art). Cell concentration and viability were assessed via Trypan blue staining using a Luna-II cell counter (Logos Biosystems). All samples had greater than 85% viability. Samples were sequenced with either single-cell RNA-seq and single-cell ATAC-seq (separately); with single-cell multiome ATAC + Gene Expression sequencing; or with all of: single-cell RNA-seq, single-cell ATAC-seq, and single-cell multiome ATAC + Gene Expression sequencing as biological replicates (**Data S2B**). We report here the single-cell RNA-seq and single-cell multiome Gene Expression portions only.

For scRNA-seq, a single-cell suspension was used for construction of scRNA libraries, which were generated with Chromium NextGEM Single-cell 3’ Gene Expression reagents (v.3.1; 10x Genomics). For scATAC-seq, a single nuclei suspension was generated following the protocol Nuclei Isolation for Single Cell ATAC Sequencing (Demonstrated protocol CG000169, Rev D; 10x Genomics). Single-cell ATAC libraries were constructed with Chromium Next GEM Single Cell ATAC reagents (v.1.1; 10x Genomics). A 10x Chromium Controller instrument was used according to the manufacturer’s instructions for both scRNA-seq and scATAC-seq, targeting a recovery of around 4,000 nuclei for each sample.

For single-cell multiome sequencing, a single nuclei suspension was generated according to protocol Nuclei Isolation for Single Cell Multiome ATAC + Gene Expression Sequencing (Demonstrated protocol CG000365, Rev D; 10x Genomics). Single-cell multiome ATAC and gene expression libraries were generated using the Chromium Next GEM Single-cell Multiome ATAC + Gene Expression reagents (10x Genomics) and a 10x Chromium Controller instrument according to the manufacturer’s instructions (User Guide G000338, Rev D), targeting a recovery of around 4,000 nuclei for each sample.

Quality control on intermediate products and final libraries was performed using the Agilent Bioanalyzer High Sensitivity Kit (Agilent Technologies). The final libraries were quantified via qPCR using KAPA Library Quantification Kits (KAPA Biosystems, #KK4824) and were sequenced on an Illumina NovaSeq6000 system or on an Illumina HiSeq 4000 system (**Data S2**) with a 1% spike-in of PhiX control library (Illumina).

### Single-cell alignment, normalization, and integration

Single-cell sequencing data were aligned to hg38. The single-omic scRNA-seq data were aligned using Cell Ranger (RRID:SCR_017344; 10x Genomics), and the multiomic data were aligned using Cell Ranger Arc (RRID:SCR_023897; 10x Genomics).

The single-cell data were analyzed using the Seurat R package v.5.1.0 (RRID:SCR_016341) (*68*). For the single-omic scRNA-seq data, cells with number of non-zero genes larger than 1,000, number of reads less than 60,000, and percentage of mitochondria reads less than 25% were retained. For the multi-omic data, only cells meet the following criteria were retained for further analysis: first, number of reads larger than 5,000 but smaller than 100,000 from the scATAC-seq data modality; second, number of reads larger than 1,000 but smaller than 100,000 from the scRNA-seq data modality; third, the percentage of mitochondria reads less than 30%. The gene expression (i.e., scRNA-seq) data were then normalized to the same library and log-transformed using the Seurat NormalizeData() function.

Samples from different patients or technologies (single or multi-omics) were integrated using Harmony (RRID:SCR_022206) (*69*) based on the gene expression data. For one sample, we obtained replicate scRNA-seq data from Stanford University (Majeti Lab; sample SU758, replicate SU758_stanford_single). For data integration, the top 2,000 most variable genes were first obtained from each sample based on the mean-adjusted variance. Then the top 2,000 genes that were consistently ranked as the most highly variable genes were used for principal component analysis (PCA). The top 30 PCs were used for integration using Harmony. Low dimensional representation of the cells and cell clusters were obtained using UMAP and Louvain clustering based on the batch-adjusted PCs from Harmony.

To annotate the cell identity of the cell clusters in the integrated single-cell data, we obtained a bulk RNA-seq dataset that contains gene expression from 13 normal hematopoietic cell types from GEO (GSE74246) (*70*). Differential gene expression analyses were applied to each pair of cell types using DESeq2 (RRID:SCR_015687) (*71*) and the union of the top 100 differential genes from each comparison was obtained. Based on this differential gene set, we calculated the Spearman’s correlation between the average gene expression from each cell cluster from the single-cell data and the quantile-normalized gene expression from the bulk data. The similarity between the cell cluster and the normal cell types were used to annotate their identities.

### Differential expression analysis

Differential expression analysis was performed using Seurat v.5.1.0 (*68*) between each AML subtype and normal samples (n=3, CD34-1, CD34-2, and GMP-1) within clusters 0, 2, and 4. Pseudobulk expression profiles were generated for each sample in each cluster using the Seurat function AggregateExpression(). Differential expression testing was performed using the FindMarkers() function with test.use = “DESeq2”. Genes were called as differentially expressed if absolute average log2 fold change ≥ 0.5 and Bonferroni-adjusted p-value ≤ 0.1.

## Calculation of gene expression mean adjusted variability

Let *y*_*ij*_ be the library-size normalized, imputed, and log2-transformed expression level for gene *i (i = 1, …, I)* and cell *j (j = 1, …, J)*. Let m_i_ and s_i_ be the mean and standard deviation of the expression level for gene *i* across all cells respectively. A LOESS regression model was fitted across all genes where *s*_*i*_ is the response variable and *m*_*i*_ is the independent variable. Let 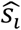 be the fitted values of the standard deviations from the regression model. The gene expression mean-adjusted variability (MAV) of gene *i, h*_*i*_, is defined as the residual of the regression model, or equivalently the difference between the observed and fitted standard deviation: 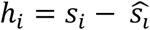.

MAV was calculated for each gene in each sample as described above. Genes were then ranked by MAV within each sample. A consensus ranking of genes for each AML subtype was generated by calculating the rank product of individual MAV ranks from each sample of the appropriate subtype. Similarly, a consensus ranking of expression level and variance for each gene in each AML subtype was generated by taking the rank product of the ranks of the expression mean or variance within each sample for the appropriate subtype. We then divided genes into quartiles of gene expression rank product, variance rank product, and MAV rank product in each AML subtype, with a higher quartile indicating a high value of the associated statistic (ex. genes in the 75-100% quartile of MAV rank product have higher MAV than genes in the 0-25% quartile). Finally, expression rank product, variance rank product, and MAV rank product were associated with values of CPEL MML and CPEL NME binned over promoter regions for visualization.

### Gene network inference

Gene network inference was performed from the single-cell gene expression data using a previously-described information-theoretic method (*52*) implemented via the Julia package NetworkInference.jl with default settings. Networks were visualized with the R package tidygraph v.1.3.1 (DOI: 10.32614/CRAN.package.tidygraph). NetworkInference.jl returns a list of every possible edge, and the confidence of each edge existing in the true network, so the top 10% of edges in the network were retained for plotting. For each gene in the network, the betweenness centrality was calculated using the function betweenness() from the R package igraph v.2.0.3 (RRID:SCR_021238). Betweenness centrality is computed as

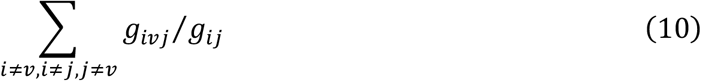

where *g*_*ij*_ is the number of shortest paths between vertices *i* and *j*, while *g*_*ivj*_ is the number of shortest paths between *i* and *j* that also pass through vertex 𝒱. Betweenness centrality measures how the gene (node) lies on paths between other nodes in the network, so a gene with the highest betweenness centrality is predicted to exert the most influence over the network.

