## Supplementary Materials for "DNA methylation stochasticity is linked to transcriptional variability and identifies convergent epigenetic disruption across genetically-defined subtypes of AML"

**The PDF file includes:**

Figures S1 to S9

**Other Supplementary Materials for this manuscript include the following:**

Data S1 to S14

**Figure S1. Overview of CPEL model**.

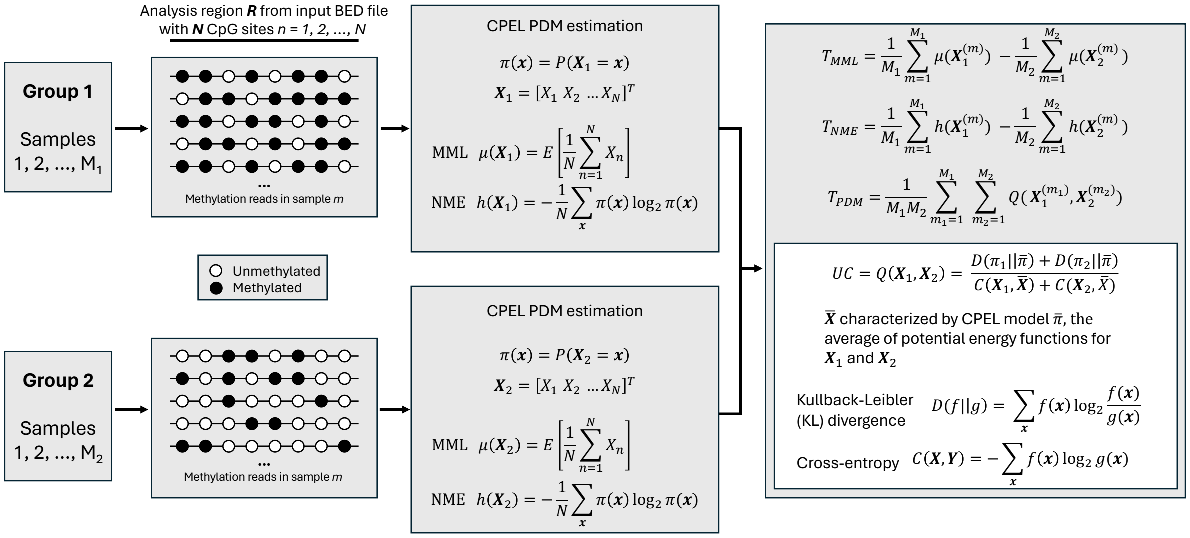


The CPEL model estimates the PDM within an analysis region *R* containing *N* CpG sites as $\pi(\boldsymbol{x}) = \mathrm{P}(\mathbf{X} = \mathbf{x})$ for a random vector $\boldsymbol{X} = [X_1\, X_2\, ..\, X_N\,]^T$ , where X = 0 represents an unmethylated CpG and X = 1 represents a methylated CpG. This PDM can be used to estimate the MML, NME, and thus the differential MML, differential NME, and difference between PDMs, quantified by the uncertainty coefficient UC.

#### Figure S2: Methylation level and entropy over genomic features.

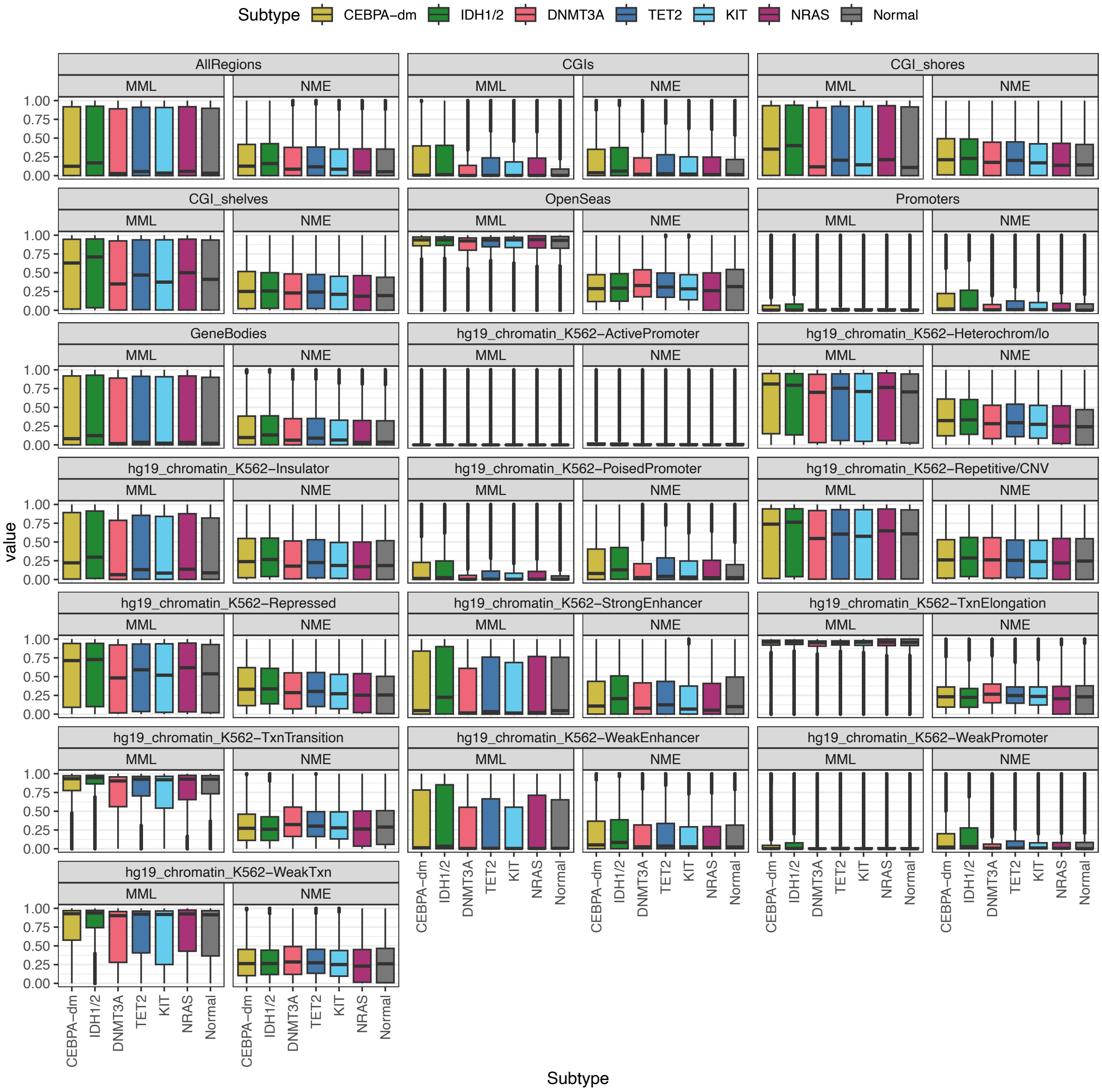


Boxplots of MML and NME as reported by CpelTdm over specified genomic features.

**Figure S3: Differential methylation stochasticity over genomic features.**

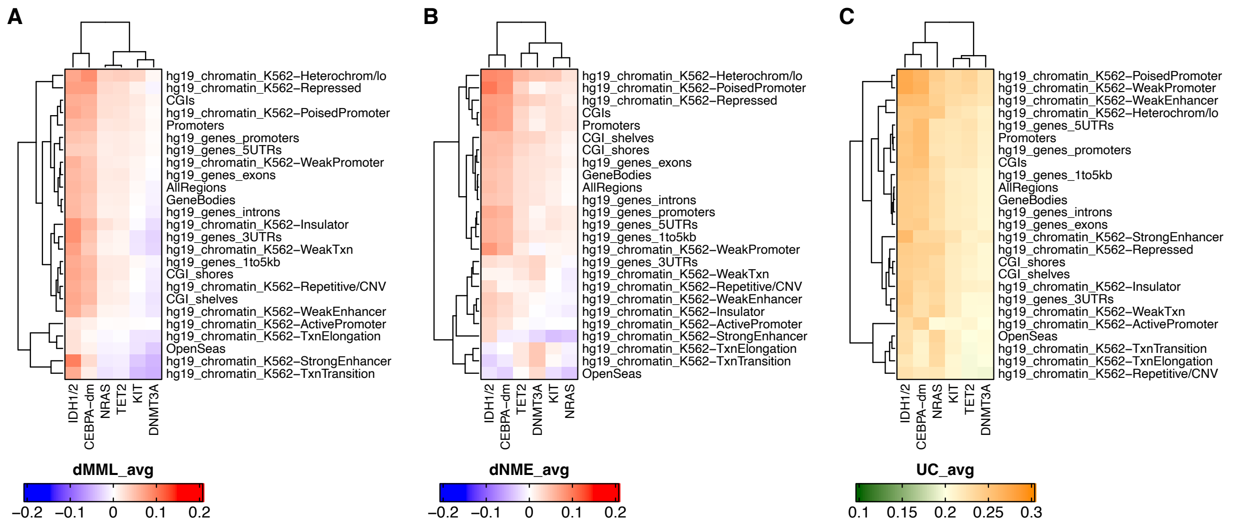


Heatmaps of (**A**) average dMML, (**B**) average dNME, and (**C**) average UC over various genomic features. CEBPA-dm and IDH1/2-mut AMLs show distinct hypermethylation, increased entropy, and high UC.

**Figure S4: Enrichment of regions with significantly discordant methylation stochasticity over genomic features.**

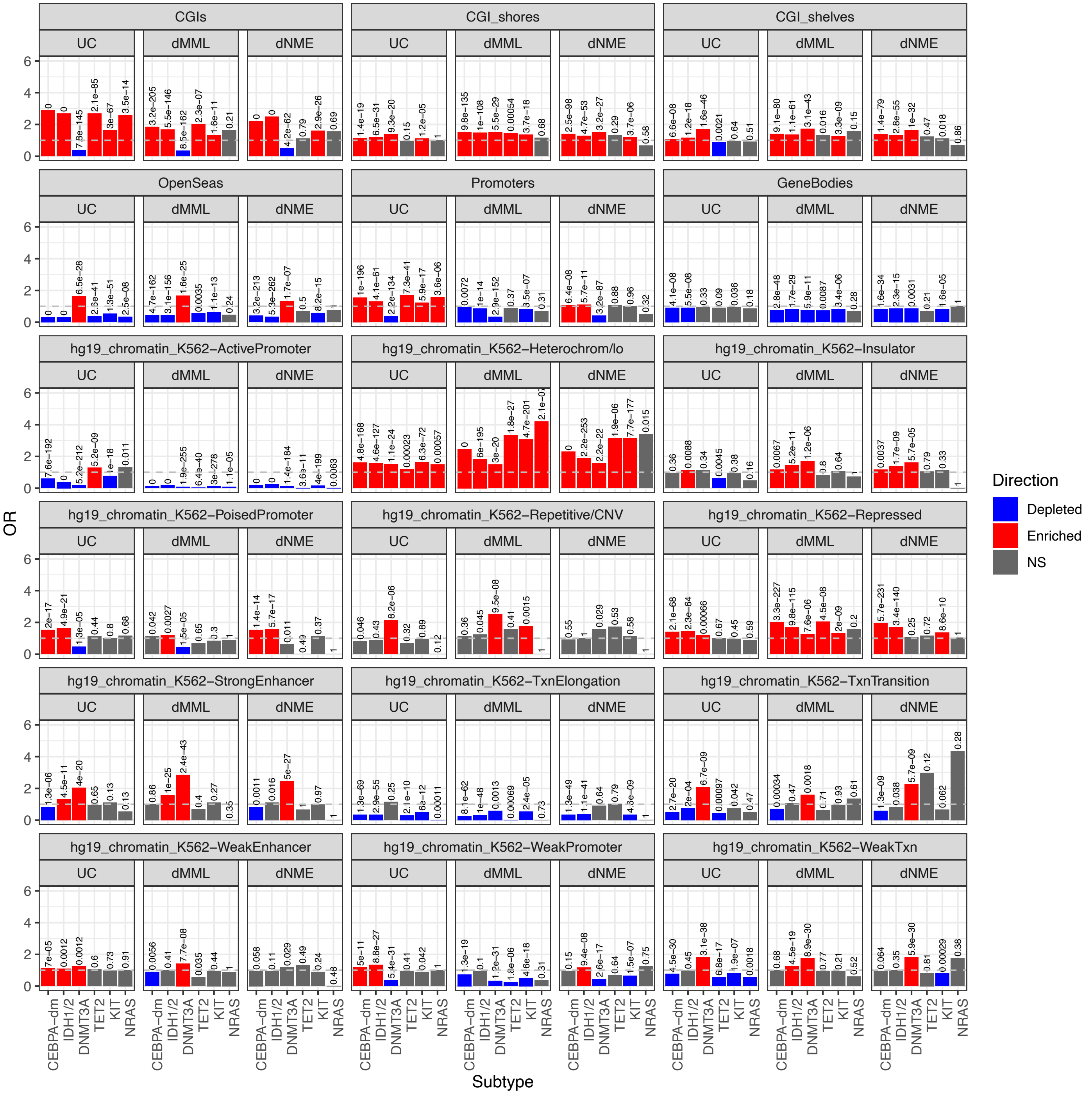


Enrichment of DMRs over genomic features was calculated using the Fisher's test (**Methods**). Color indicates direction (depleted = blue, odds ratio < 1 and Benjamini-Hochberg adjusted p-value ≤ 0.01; enriched = red, odds ratio > 1 and Benjamini-Hochberg adjusted p-value ≤ 0.01). The number above each bar indicates the BH-adjusted p-value for the given enrichment test.

**Figure S5: Transcription factor binding motif enrichment of regions with significantly discordant methylation stochasticity.**

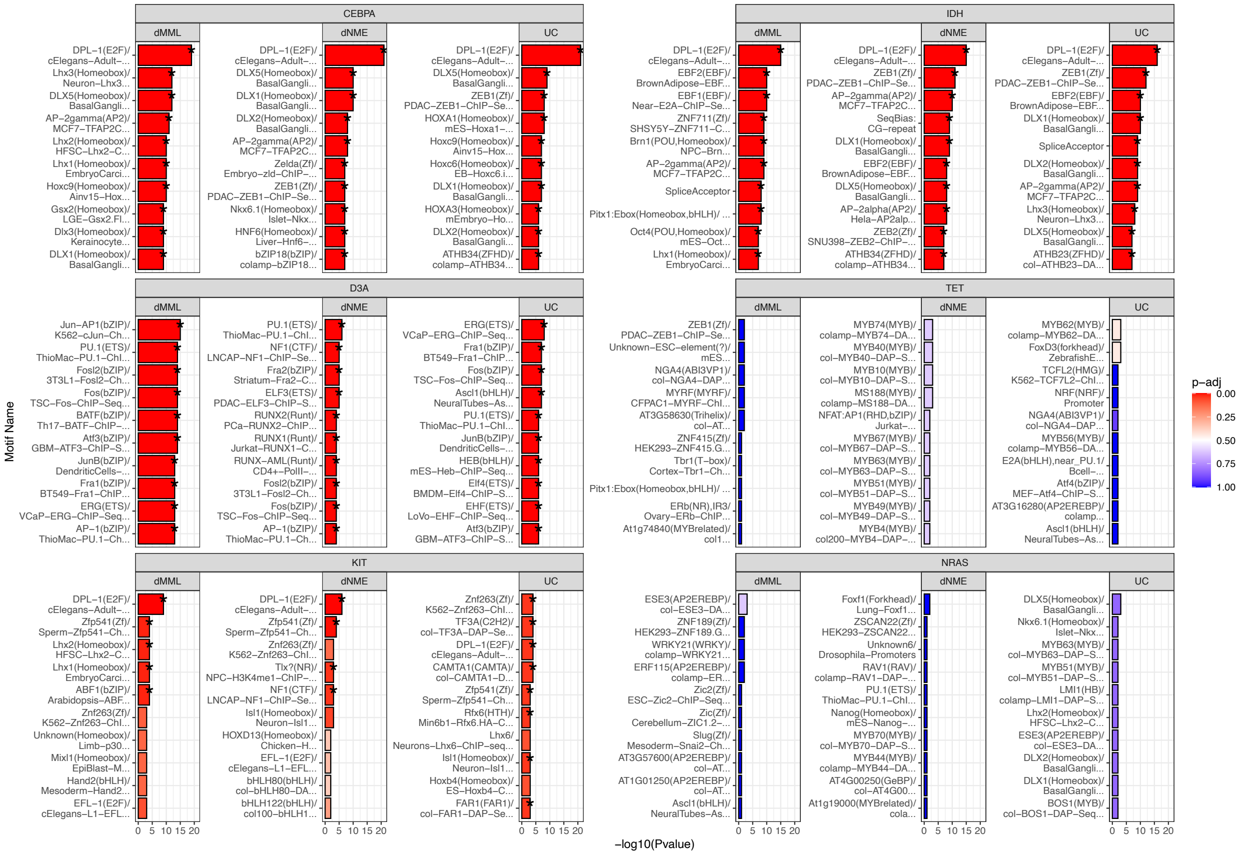


HOMER known transcription factor motif enrichment analysis results for dMML DMRs, dNME DMRs, and UC DMRs. DMRs in CEBPA-dm and IDH1/2-mut AMLs are enriched in homeobox and AP-2 family binding motifs, while DMRs in DNMT3A-mut AMLs are enriched in ETS and RUNT family motifs. Color indicates adjusted p-value. *: adjusted p-value ≤ 0.05.

**Figure S6: Discordant methylation stochasticity in AML localizes to leukemia signatures and regulators of oncogenesis.**

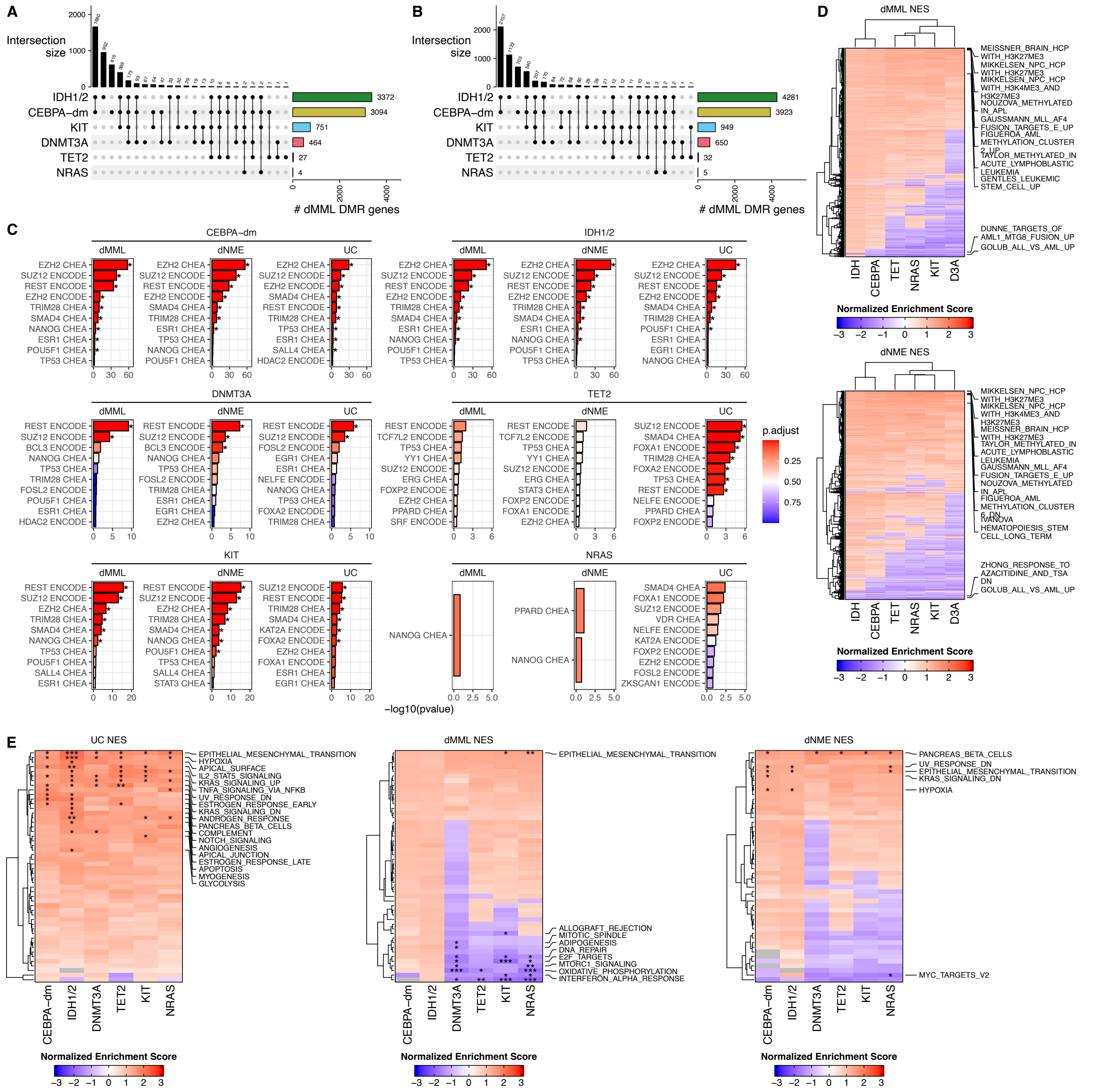


Overlap of genes associated with dMML DMRs **(A)** and dNME DMRs **(B)** in AML subtypes. **(C)** Enrichment of genes with dMML, dNME, and UC DMRs in transcription factor target gene sets by over-representation analysis (**Methods**). *: BH adjusted p-value ≤ 0.05. **(D)** Gene set enrichment analysis of genes ranked by dMML (top) or dNME (bottom) over the MSigDB C2 gene set library. Selected gene sets related to histone marks, leukemia, and hematopoietic signatures are annotated. **(E)** Gene set enrichment analysis of genes ranked by UC (left), dMML (middle), or dNME (right) over the MSigDB Curated Hallmark gene set library. *: BH adjusted p-value ≤ 0.05; **: adjusted p-value ≤ 1e-3; ***: adjusted p-value ≤ 1e-4. For UC, positive NES = high UC (highly discordant methylation stochasticity) and negative NES = low UC (lowly discordant methylation stochasticity); for dNME, positive NES = increased NME in AML and negative NES = decreased NME in AML; for dMML, positive NES = increased MML in AML and negative NES = decreased MML in AML.

**Figure S7: Single-cell RNA-seq analysis identifies dominant cell type identities in AML.**

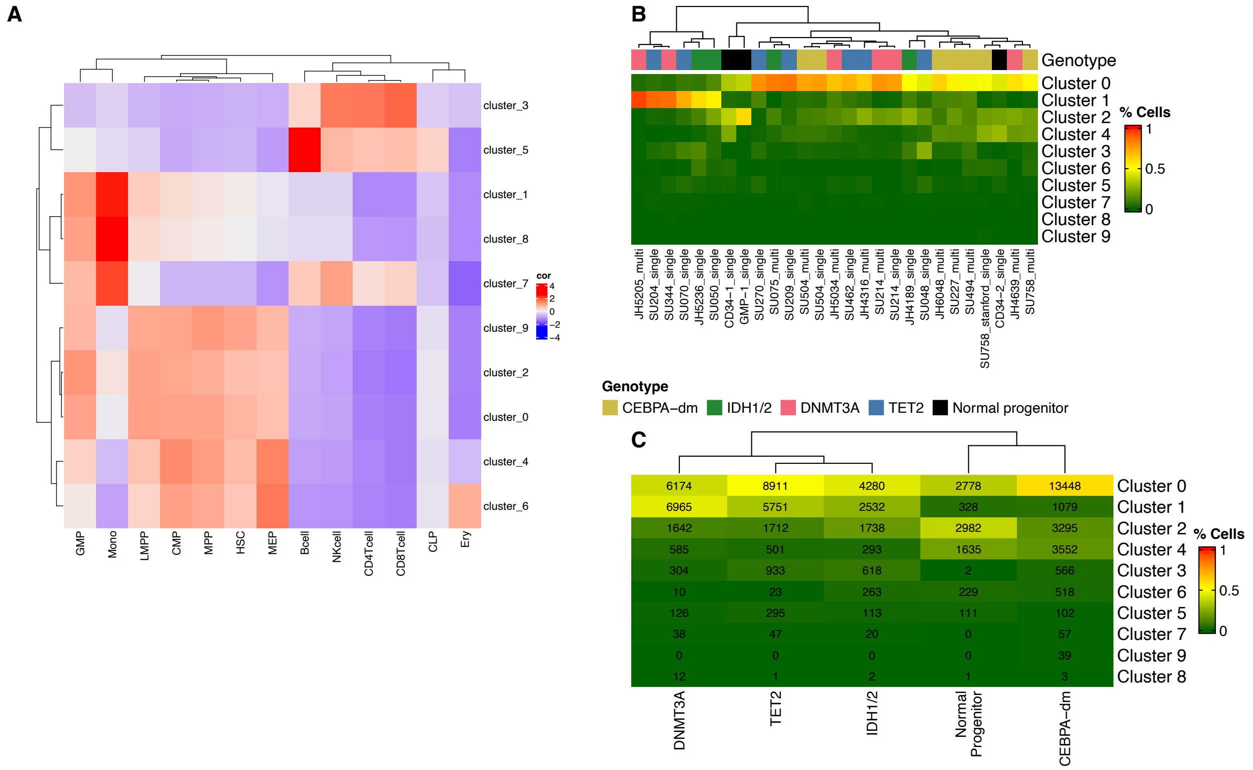


**(A)** Heatmap of Spearman correlation between cluster expression profiles (rows) and defined hematopoietic gene expression profiles (**Methods**). **(B)** Proportion of cells in each sample (columns) in each cluster (rows). The genotype for each sample is indicated by the color above each column. **(C)** Proportion of cells in each subtype (AML and normal) (columns) in each cluster (rows). Annotations indicate the number of cells in each cluster-subtype designation.

#### Figure S8: Transcriptional dysregulation recapitulates leukemia signatures.

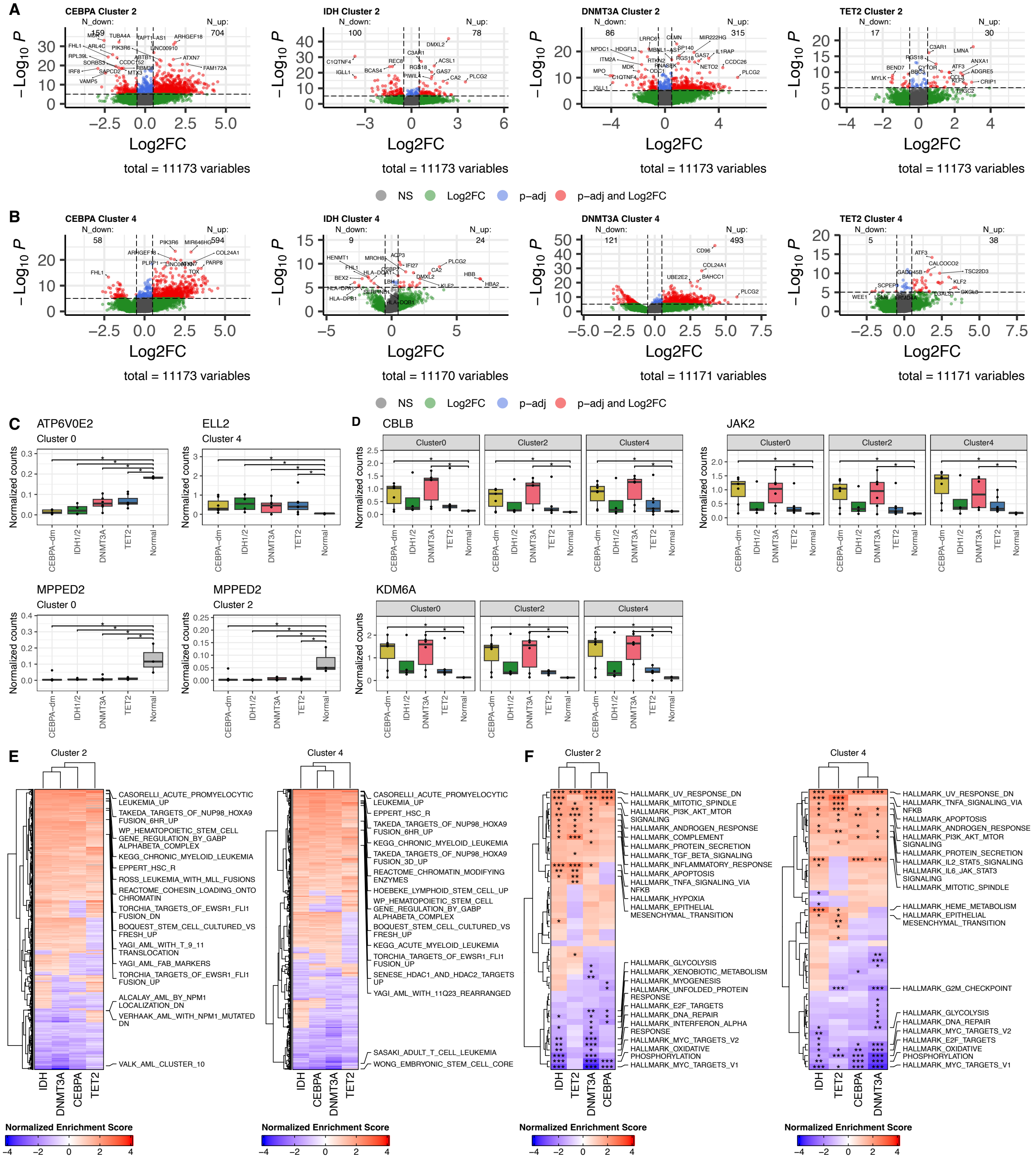

Volcano plots of differentially expressed genes in Cluster 0 for CEBPA-dm AMLs (left), IDH1/2-mutant AMLs (middle-left), DNMT3A-mutant AMLs (middle-right), and TET2-mutant AMLs (right) versus normal progenitors in **(A)** cluster 2 and **(B)** cluster 4. Genes were called as differentially expressed if absolute average log2 fold change ≥ 0.5 and Bonferroni-adjusted p-value ≤ 0.1. **(C)** Boxplots of cluster-level pseudobulk expression profiles for genes differentially expressed in all AML versus normal comparisons (within a given cluster). Panel titles indicate the corresponding gene and cluster. *: Bonferroni adjusted p-value ≤ 0.05. **(D)** Boxplots of cluster-level pseudobulk expression profiles for representative AML genetic drivers differentially expressed in AML versus normal comparisons (within a given cluster). Panel titles indicate the corresponding gene and cluster. *: Bonferroni adjusted p-value ≤ 0.05. **(E)** Gene set enrichment analysis over the MSigDB C2 gene set library of genes ranked by average log2 fold change (versus normal samples) in clusters 2 (left) and 4 (right). Selected gene sets related to histone marks, leukemia, and hematopoietic signatures are annotated. **(F)** Gene set enrichment analysis over the MSigDB Curated Hallmark gene set library of genes ranked by average log2 fold change (versus normal samples) in clusters 2 (left) and 4 (right). *: Benjamini-Hochberg adjusted p-value ≤ 0.05; **: adjusted p-value ≤ 1e-3; ***: adjusted p-value ≤ 1e-4.

### Figure S9. Methylation stochasticity relates to gene expression across AML subtypes.

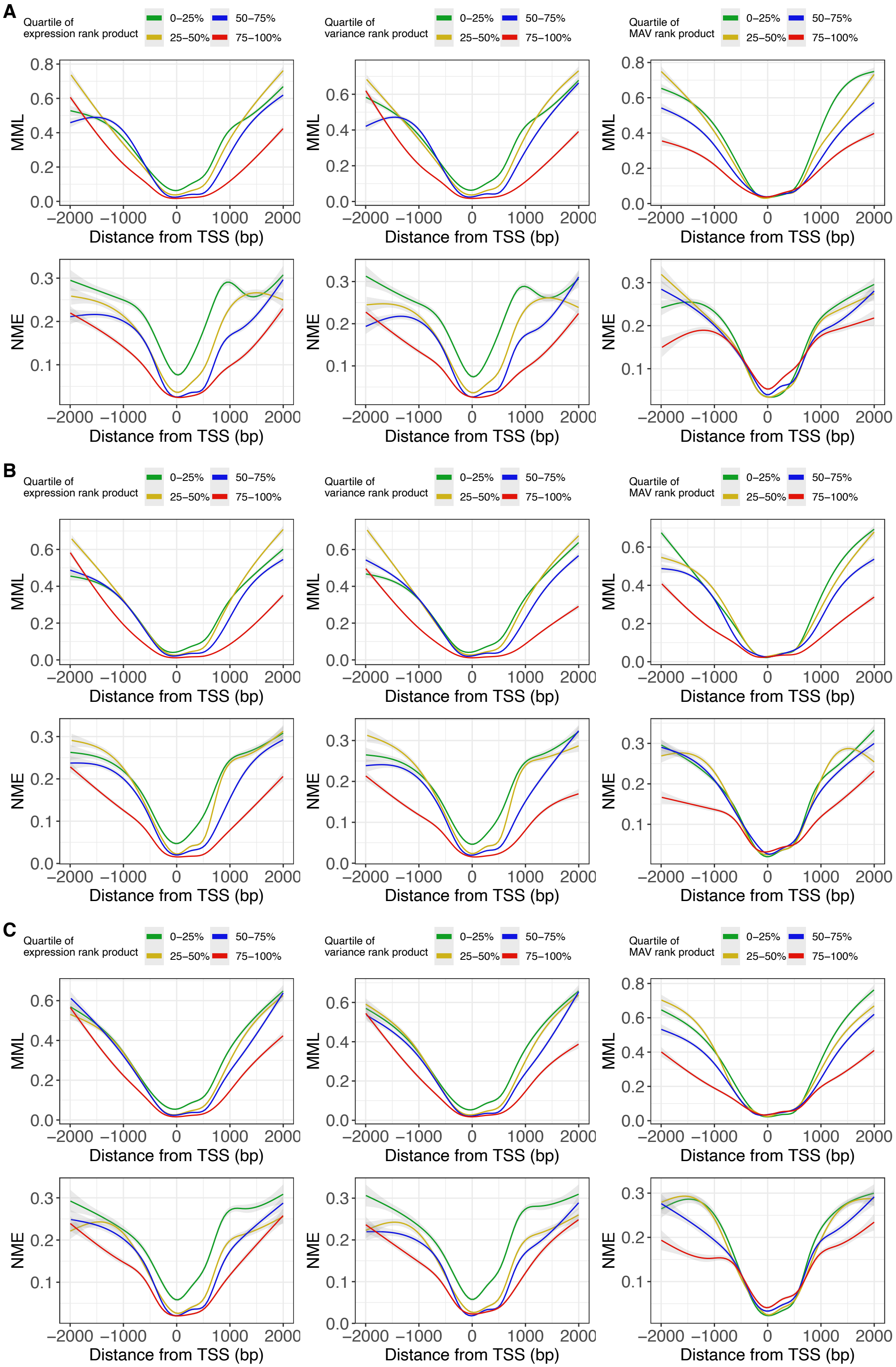

Relationship between MML or NME and gene expression mean (left), variance (middle), or variability (MAV; right) in CEBPA-dm (**A**), DNMT3A-mut (**B**), and TET2-mut (**C**) AMLs, as a function of distance from the TSS. Lower methylation level is associated with higher expression level (top-left), while lower expression levels are associated with lower levels of NME (bottom-left). Higher expression variance is associated with lower MML and NME, but this may be driven by the relationship between mean expression and expression variance (**Methods**). Mean-adjusted expression variability is associated with higher NME near the TSS, connecting methylation stochasticity to expression variability.

**Data S1. Results for CPEL runs defining analysis regions using various coverage strategies.**

The CpelTdm.jl package was applied to ERRBS data over analysis regions defined by subsetting CpGs using various coverage strategies (**Methods**). The number of DMRs at different q-value cutoffs are reported here. The columns are as follows:

- Coverage strategy: The coverage strategy implemented
- Comparison (vs normal): AML subtype vs normal comparison
- NumberEvaluatedRegions: Number of analysis regions evaluated
- kbEvalRegions: kb covered by evaluated analysis regions
- AverageCpGsperRegion: Average number of CpGs within analysis regions
- numberUC_DMRsq02: Number of analysis regions with significant UC ($q \leq 0.2$)
- kbUC_DMRsq02: kb of analysis regions with significant UC ($q \leq 0.2$)
- numberdMML_DMRsq02: Number of analysis regions with significant dMML ($q \leq 0.2$)
- kbdMML_DMRsq02: kb of analysis regions with significant dMML ($q \leq 0.2$)
- numberdNME_DMRsq02: Number of analysis regions with significant dNME ($q \leq 0.2$)
- kbdNME_DMRsq02: kb of analysis regions with significant dNME ($q \leq 0.2$)
- numberUCDMRsq01: Number of analysis regions with significant UC ($q \leq 0.1$)
- kbUCDMRsq01: kb of analysis regions with significant UC ($q \leq 0.1$)
- numberdMML_DMRsq01: Number of analysis regions with significant dMML ($q \leq 0.1$)
- kbdMML_DMRsq01: kb of analysis regions with significant dMML ($q \leq 0.1$)
- numberdNME_DMRsq01: Number of analysis regions with significant dNME ($q \leq 0.1$)
- kbdNME_DMRsq01: kb of analysis regions with significant dNME ($q \leq 0.1$)

**Data S2. Results for CPEL runs defining analysis regions using various coverage strategies.**

**Sheet A**

Mutational profiles for the 43 AML samples used for methylation stochasticity analysis (ERRBS data). Samples were downloaded from publicly available data (**Methods**). Columns are as follows:

- Sample: AML sample name
- Run: SRA accession
- GEOaccession: GEO accession
- AML subtype: AML subtype assigned based on mutational profile (CEBPA-dm, IDH1/2, DNMT3A, TET2, KIT, or NRAS)
- Age: Age of patient at point of sample collection
- Gender: Patient gender
- FAB: AML sample FAB classification
- Cytogenetics: Cytogenetic profile of sample
- Columns I-AX indicate presence or absence of the mutation in the column title; ie: a sample with "Pos" in the column "DNMT3A" has a mutation in DNMT3A.

**Sheet B**

Mutational profiles and sequencing information for the 42 AML samples used for single-cell RNA/ATAC-seq analysis. Samples were obtained from the Johns Hopkins Hospital or Stanford University biobank (**Methods**). Columns are as follows:

- Sample: AML sample name
- AML subtype: AML subtype assigned based on mutational profile (CEBPA-dm, IDH1/2, DNMT3A, TET2, mixed/other mutations, or normal)
- Sequencing_type: Type of sequencing performed on sample (scRNA-seq and scATAC-seq, and/or sc-Multiome ATAC + Gene Expression sequencing). Only scRNA-seq and gene expression data are reported in this paper.
- Sequencing_system: Name of Illumina system used for sequencing
- Sample_source: Source of sample
- Dx: Patient diagnosis
- Cytogenetics: Cytogenetic profile of sample
- Columns F-AA indicate presence or absence of the mutation in the column title; ie: a sample with "Pos" in the column "DNMT3A" has a mutation in DNMT3A.

**Data S3**: **CPEL analysis regions with associated CPEL methylation stochasticity statistics.**

The CpelTdm.jl package was applied to each AML subtype versus normal comparison (**Methods**). The CPEL statistics characterizing methylation stochasticity (dMML, dNME, and UC) are reported here for each AML versus normal comparison, using the analysis regions generated by requiring CpGs to have 10x coverage in all samples, along with any genes associated with a given analysis region.

Sheet name indicates the AML vs normal comparison (ex. sheet 10xCov_allSamps_CEBPAdm reports statistics from the CEBPA-dm vs. normal comparison).

Columns are as follows:

- Region: hg19 coordinates of analysis region
- dMML_score: dMML value
- dMML_p.val: p-value associated with MML comparison
- dMML_p.adj: adjusted p-value associated with MML comparison
- dNME_score: dNME value
- dNME_p.val: p-value associated with NME comparison
- dNME_p.adj: adjusted p-value associated with NME comparison
- UC_score: UC value
- UC_p.val: p-value associated with UC/PDM comparison
- UC_p.adj: adjusted p-value associated with UC/PDM comparison
- UC_dmr_ind: Indicator of UC DMR status ($q \leq 0.2$), UC_dmr or NS (not significant)
- dNME_dmr_ind: Indicator of dNME DMR status ($q \leq 0.2$), dNME_dmr or NS (not significant)
- dMML_dmr_ind: Indicator of dMML DMR status ($q \leq 0.2$), dMML_dmr or NS (not significant)
- gene_symbol: Gene symbol of gene(s) containing the analysis region in the promoter (**Methods**). If the analysis region is present in the promoter of multiple genes, gene names are separated by a comma (“,”).

**Data S4**: **HOMER knownMotif transcription factor binding motif enrichment analysis results for dMML, dNME, and UC DMRs identified by CPEL analysis.**

HOMER motif analysis was run on genomic regions containing dMML, dNME, and UC DMRs from CPEL analysis (**Methods**). Reported here are results from HOMER run on DMRs identified by CPEL with analysis regions defined by requiring CpGs to have 10x coverage in all samples (**Methods**).

Sheet names indicate the AML subtype versus normal comparison and the CPEL statistic used to identify DMRs (ex. Sheet cebpa_dMML contains results from HOMER run on dMML DMRs identified in a CEBPA-dm vs normal CPEL comparison).

Columns are as follows, reported without modification from the HOMER knownMotif output:

- MotifName: Name of HOMER known motif
- Consensus: Consensus sequence of binding motif
- Pvalue: P-value associated with motif enrichment
- LogPvalue: Log of the p-value associated with motif enrichment
- Qvalue: Adjusted p-value associated with motif enrichment
- # of Target Sequences with Motif: Number input regions containing the HOMER motif
- % of Target Sequences with Motif: Percent input regions containing the HOMER motif
- # of Background Sequences with Motif: Number background regions containing the HOMER motif
- % of Background Sequences with Motif: Percent background regions containing the HOMER motif

**Data S5: Intersection of genes associated with UC DMRs in CEBPA-dm and IDH1/2-mut AMLs.**

DMRs were mapped to associated genes based on the presence in the promoter region (**Methods**, **Table S2**). Here, we report the intersection of genes associated with UC DMRs in CEBPA-dm and IDH1/2-mut AMLs and genes associated with UC DMRs in all AML subtypes, using DMRs identified by CPEL with analysis regions defined by requiring CpGs to have 10x coverage in all samples (**Methods**).

**Data S6**: **Over-representation analysis of DMR-associated genes over transcription factor target gene sets.**

Over-representation analysis (ORA) for genes associated with dMML, dNME, and UC DMRs was performed with the clusterProfiler package over the "ENCODE_and_ChEA_Consensus_TFs_from_ChIP-X" gene set library downloaded from Enrichr (**Methods**). The results of the ORA are presented here for each AML subtype versus normal comparison and each CPEL statistic.

Sheet names indicate the AML subtype versus normal comparison and the CPEL statistic used to identify DMR-associated genes (ex. Sheet cebpa_dMMLdmrGenes contains results from clusterProfiler ORA run on genes with dMML DMRs identified in a CEBPA-dm vs normal CPEL comparison).

Column names are as follows:

- ID: Gene set ID
- Description: Gene set description
- GeneRatio: clusterProfiler gene ratio statistic, calculated as k/n where k is the overlap between the input gene list and the gene set, and n is the total number of input genes
- BgRatio: clusterProfiler background ratio statistic, calculated as M/N where M is the overlap between the background gene list and the gene set, and N is the total number of background genes
- pvalue: clusterProfiler p-value
- p.adjust: clusterProfiler Benjamini-Hochberg adjusted p-value
- qvalue: clusterProfiler q-value, estimated using the method of the R qvalue package (10.18129/B9.bioc.qvalue)
- geneID: List of input genes overlapping the query gene set, separated by "/"
- Count: Number of input genes overlapping the query gene set

**Data S7: Gene set enrichment analysis of genes ranked by CPEL statistics over MSigDB C2 gene set collection.**

Gene set enrichment analysis (GSEA) for genes ranked by dMML, dNME, and UC was performed with the clusterProfiler package over the MSigDB C2 gene set collection (**Methods**). The results of the GSEA are presented here for each AML subtype versus normal comparison and each CPEL statistic.

Sheet names indicate the AML subtype versus normal comparison and the CPEL statistic used to identify DMR-associated genes (ex. Sheet cebpa_dMML contains results from clusterProfiler GSEA run on genes ranked by the strength of the dMML from a CEBPA-dm vs normal CPEL comparison).

Column names are as follows:

- ID: Gene set ID
- Description: Gene set description
- setSize: Number of genes in the gene set, subset by genes in the background set
- enrichmentScore: GSEA enrichment score calculated via leading-edge analysis
- NES: GSEA normalized enrichment score
- pvalue: clusterProfiler p-value
- p.adjust: clusterProfiler Benjamini-Hochberg adjusted p-value
- qvalue: clusterProfiler q-value, estimated using the method of the R qvalue package (10.18129/B9.bioc.qvalue)
- Rank: clusterProfiler GSEA rank output
- leading_edge: Leading edge analysis statistics
- core_enrichment: List of input genes overlapping the gene set, separated by "/"

**Data S8: Gene set enrichment analysis of genes ranked by CPEL statistics over MSigDB Hallmark gene set collection.**

Gene set enrichment analysis (GSEA) for genes ranked by dMML, dNME, and UC was performed with the clusterProfiler package over the MSigDB Hallmark gene set collection (**Methods**). The results of the GSEA are presented here for each AML subtype versus normal comparison and each CPEL statistic.

Sheet names indicate the AML subtype versus normal comparison and the CPEL statistic used to identify DMR-associated genes (ex. Sheet cebpa_dMML contains results from clusterProfiler GSEA run on genes ranked by the strength of the dMML from a CEBPA-dm vs normal CPEL comparison).

Column names are as follows:

- ID: Gene set ID
- Description: Gene set description
- setSize: Number of genes in the gene set, subset by genes in the background set
- enrichmentScore: GSEA enrichment score calculated via leading-edge analysis
- NES: GSEA normalized enrichment score
- pvalue: clusterProfiler p-value
- p.adjust: clusterProfiler Benjamini-Hochberg adjusted p-value
- qvalue: clusterProfiler q-value, estimated using the method of the R qvalue package (10.18129/B9.bioc.qvalue)
- Rank: clusterProfiler GSEA rank output
- leading_edge: Leading edge analysis statistics
- core_enrichment: List of input genes overlapping the gene set, separated by "/"

**Data S9: Differential expression analysis of AML samples versus normal progenitors.**

Differential expression analysis was performed after pseudobulking for each sample within clusters 0, 2, and 4 (**Methods**). The results of the analysis are presented here for each AML subtype versus normal comparison and each analyzed cluster.

Sheet names indicate the AML subtype versus normal comparison and the cluster (ex. Sheet DNMT3A_Cluster0 contains differential expression results from the DNMT3A versus normal, within cluster 0).

Column names are as follows:

- p_val: p-value returned by Seurat DESeq2 analysis (**Methods**)
- avg_log2FC: Log2 fold change between AML and normal. Negative log2FC indicates downregulation in AML; positive log2FC indicates upregulation in AML.
- pct.1: Percent of AML samples expressing the given gene.
- pct.2: Percent of normal samples expressing the given gene.
- p_val_adj: Bonferonni-adjusted p-value
- Gene: Gene symbol

**Data S10: Differentially expressed genetic drivers of AML.**

We generated a list of known genetic drivers of AML (Papaemmanuil et al., 2016). Here, we report the set of these genetic drivers that were identified as DEGs in AML versus normal comparisons (**Methods**).

Column names are as follows:

- AML genetic driver: The gene symbol of the known genetic driver of AML
- Subtype: The AML subtype in which the genetic driver is a DEG (versus normal)
- Direction: Differential expression direction (up = upregulated in AML, down = downregulated in AML)
- Cluster: The cluster in which the genetic driver is a DEG

**Data S11: Over-representation analysis of differentially expressed genes over transcription factor target gene sets.**

Over-representation analysis (ORA) for DEGs in clusters 0, 2, and 4 was performed with the clusterProfiler package over the "ENCODE_and_ChEA_Consensus_TFs_from_ChIP-X" gene set library downloaded from Enrichr (**Methods**). The results of the ORA are presented here for each AML subtype versus normal comparison and each cluster. Additionally, enrichment for upregulated genes and downregulated genes was calculated separately.

Sheet names indicate the AML subtype versus normal comparison, the cluster in which the comparison was run, and the direction of the DEGs used for ORA (ex. Sheet CEBPA_Cluster0_allDEGs contains results from clusterProfiler ORA run on all DEGs identified in the CEBPA versus normal comparison within cluster 0; and sheet CEBPA_Cluster0_downregulated contains results from clusterProfiler ORA run on only downregulated genes identified in the CEBPA versus normal comparison within cluster 0).

Column names are as follows:

- ID: Gene set ID
- Description: Gene set description
- GeneRatio: clusterProfiler gene ratio statistic, calculated as k/n where k is the overlap between the input gene list and the gene set, and n is the total number of input genes
- BgRatio: clusterProfiler background ratio statistic, calculated as M/N where M is the overlap between the background gene list and the gene set, and N is the total number of background genes
- pvalue: clusterProfiler p-value
- p.adjust: clusterProfiler Benjamini-Hochberg adjusted p-value
- qvalue: clusterProfiler q-value, estimated using the method of the R qvalue package (10.18129/B9.bioc.qvalue)
- geneID: List of input genes overlapping the query gene set, separated by "/"
- Count: Number of input genes overlapping the query gene set

**Data S12: Gene set enrichment analysis of genes ranked by log2 fold change over MSigDB C2 gene set collection.**

Gene set enrichment analysis (GSEA) for genes ranked by log2 fold change (versus normal samples) was performed with the clusterProfiler package over the MSigDB C2 gene set collection (**Methods**), where positive log2 fold change = upregulated in AML and negative log2 fold change = downregulated in AML. The results of the GSEA are presented here for each AML subtype versus normal comparison in each of clusters 0, 2, and 4.

Sheet names indicate the AML subtype versus normal comparison and the cluster in which the differential expression testing was run (ex. Sheet CEBPA_Cluster0 contains results from clusterProfiler GSEA run on genes ranked by the log2 fold change from the CEBPA-dm vs normal comparison within cluster 0).

Column names are as follows:

- ID: Gene set ID
- Description: Gene set description
- setSize: Number of genes in the gene set, subset by genes in the background set
- enrichmentScore: GSEA enrichment score calculated via leading-edge analysis
- NES: GSEA normalized enrichment score
- pvalue: clusterProfiler p-value
- p.adjust: clusterProfiler Benjamini-Hochberg adjusted p-value
- qvalue: clusterProfiler q-value, estimated using the method of the R qvalue package (10.18129/B9.bioc.qvalue)
- Rank: clusterProfiler GSEA rank output
- leading_edge: Leading edge analysis statistics
- core_enrichment: List of input genes overlapping the gene set, separated by “/”

**Data S13: Gene set enrichment analysis of genes ranked by log2 fold change over MSigDB Hallmark gene set collection.**

Gene set enrichment analysis (GSEA) for genes ranked by log2 fold change (versus normal samples) was performed with the clusterProfiler package over the MSigDB Hallmark gene set collection (**Methods**), where positive log2 fold change = upregulated in AML and negative log2 fold change = downregulated in AML. The results of the GSEA are presented here for each AML subtype versus normal comparison in each of clusters 0, 2, and 4.

Sheet names indicate the AML subtype versus normal comparison and the cluster in which the differential expression testing was run (ex. Sheet CEBPA_Cluster0 contains results from clusterProfiler GSEA run on genes ranked by the log2 fold change from the CEBPA-dm vs normal comparison within cluster 0).

Column names are as follows:

- ID: Gene set ID
- Description: Gene set description
- setSize: Number of genes in the gene set, subset by genes in the background set
- enrichmentScore: GSEA enrichment score calculated via leading-edge analysis
- NES: GSEA normalized enrichment score
- pvalue: clusterProfiler p-value
- p.adjust: clusterProfiler Benjamini-Hochberg adjusted p-value
- qvalue: clusterProfiler q-value, estimated using the method of the R qvalue package (10.18129/B9.bioc.qvalue)
- Rank: clusterProfiler GSEA rank output
- leading_edge: Leading edge analysis statistics
- core_enrichment: List of input genes overlapping the gene set, separated by "/"

**Data S14: Gene expression mean-adjusted variability, variance, and mean in AML subtypes**

Mean-adjusted variability (MAV), variance, and mean expression was calculated for each gene in each sample, and genes were ranked by each statistic (**Methods**). A consensus ranking for each gene in each AML subtype and normal samples was calculated by taking the rank product of the ranks of MAV, variance, or mean expression in each sample of a given AML subtype. Reported here are the rank product and averages of each statistic, as well as the value and rank in each individual sample. For normal samples, PBMCs were excluded from rank product and average calculations.

Sheet names correspond to subtypes: Normal, DNMT3A, IDH, TET2, and CEBPA.

Columns are as follows:

- gene_symbol
- RankProduct_MAV: Rank product of the gene rankings by expression MAV within each subtype
- RankProduct_exprmean: Rank product of the gene rankings by mean expression level within each subtype
- RankProduct_var: Rank product of the gene rankings by expression variance within each subtype
- average_exprmean: average of the mean expression levels from each sample within each subtype
- average_MAV: average of the expression MAV from each sample within each subtype
- average_var: average of the expression variance from each sample within each subtype

Columns with sample names in the column header:

- exprmean_(sample name): Mean expression level for the given gene within the sample indicated by the sample name in the column title
- exprmean_rank_(sample name): Rank of the mean expression level for the given gene within the sample indicated by the sample name in the column title. A high rank (small value) indicates the gene has high expression in the sample.
- MAV_(sample name): MAV for the given gene within the sample indicated by the sample name in the column title
- MAV_rank_(sample name): Rank of the MAV for the given gene within the sample indicated by the sample name in the column title. A high rank (small value) indicates the gene has high MAV in the sample.
- var_(sample name): Variance for the given gene within the sample indicated by the sample name in the column title
- var_rank_(sample name): Rank of the variance for the given gene within the sample indicated by the sample name in the column title. A high rank (small value) indicates the gene has high variance in the sample.
